# Mapping the PTEN Mutation Landscape: Structural and Functional Drivers of Lung Cancer

**DOI:** 10.1101/2024.10.06.616856

**Authors:** Mohammad Uzzal Hossain, Mohammad Nazmus Sakib, A.B.Z. Naimur Rahman, SM Sajid Hasan, Nazia Hassan Nisha, Arittra Bhattacharjee, Zeshan Mahmud Chowdhury, Ishtiaque Ahammad, Keshob Chandra Das, Mohammad Shahedur Rahman, Md. Salimullah

## Abstract

Lung cancer is the predominant form of cancer globally, arising from the dysfunction of genetic mutations. Although PTEN mutation is crucial in the aetiology of lung cancer, the mapping of these major drivers has to be determined. We leverage computational algorithms on 43,855 SNPs of PTEN to discover the mutational impact contributing to lung cancer. Fifteen variations were identified as detrimental, and no pertinent studies have previously addressed their structural and functional aspects. Notably, seven variations were identified as the most significant contributors to lethal effects in functional aberration, as demonstrated by the computational assessment. Subsequently, molecular simulation elucidated the structural instability associated with these alterations. Furthermore, drug binding experiments at the mutational site corroborated the destabilization experiments by demonstrating the conformational alteration of the structure, resulting in varied amino acid interactions. In summary, the present study elucidates the influence of mutations in PTEN structure on its functional architecture.

## Introduction

Lung cancer is the most frequent type of cancer globally. GLOBOCAN estimates 2.4 million new cases and 1.8 million deaths from lung cancer in 2022, an increase from previous years ^1^ There are multiple risk factors of lung cancer, tobacco smoking is one of the predominant risk factors that are found in 80-90% of lung cancer^2^. Tumors from lung cancer can result from genetic abnormalities caused by a variety of carcinogenic factors ^3^. Non-small cell lung cancers (NSCLC) and small cell lung cancer (SCLC) are the two histologic classifications for these cancers. Statistics show that between 80% and 85% of all lung cancer cases are NSCLC, with large cell carcinomas accounting for 10% to 15%, adenocarcinomas (AC) for 40%, and squamous cell carcinomas (SQCC) for 25% to 30%^4^. The most common mutations in lung cancer in NSCLC are discovered in the tumor suppressor TP53 gene. Mutations that cause loss of function in tumor suppressor genes which are frequently found in adenocarcinoma include LKB1/STK11, NF1, CDKN2A, SMARCA4, and KEAP1. Tumor suppressors that are inactivated in SQCCs differ from those in AC and partially overlap with it. Most tumors contain TP53, while other tumor suppressors that are inactivated include CDKN2A, PTEN, KEAP1, MLL2, HLA-A, NFE2L2, NOTCH1, and RB1^5^. Therefore, it is essential to identify the mutations responsible for lung cancer to further guidance in diagnosis and medication.

With the advancement of genetic technology and the Human Genome Project, next-generation sequencing has presented us with a completely new understanding of genetic variants in various lung cancer subtypes ^6^. All forms of somatically acquired mutations in lung cancer may now be cataloged thanks to extensive sequencing methods like DNA whole exome sequencing^7^. These mutations include base substitutions, insertions and deletions, copy number alterations, and genomic rearrangements. Unprecedented insights into the pathophysiology of human lung cancer have been offered by the application of high-coverage sequencing technology. Specially, a large number of Single Nucleotide Polymorphisms (SNP) has been identified which are available in the databases. Functional changes are more likely to result from SNPs in the regulatory regions or coding regions of genes (cSNPs) than from SNPs elsewhere. SNPs are the most prevalent type of genetic mutation, accounting for over 90% of the variation in the human genome^8^. The human genome has a large number of genetic polymorphisms, therefore studying each one in depth is required to determine its relevance. Further, each SNP presented in the database may not pose the threat for a disease and sequencing technologies are not enough to understand their contribution in generating a specific disease and development of innovative drugs for individuals^9^

The introduction of structural assessment via deleterious variant identification provides a great chance to overcome the challenging task of testing every SNPs in wet lab settings. As a result, employing specific procedures such as the *in-silico* technique is an effective means of identifying which SNPs are detrimental^10^. Furthermore, integrative analysis of different algorithms enhances the accuracy of the projected consequences of specific mutations. In addition, state-of-the-art methods like molecular dynamics modeling enable accurate evaluation of modifications to protein characteristics, such as structure, chemical composition, and interactions, in a simulated setting. The potential significance of mutation in comprehending the molecular pathways of different diseases is revealed by recent studies on the nsSNPs employing computational techniques ^11^

Hence, aiming to identify the damaging effect to protein structures and gene regulation, we conduct experiments using computational tools that are based on different algorithms and technologies which help to identify the potential impacts of alteration. These tools take different factors into consideration including damaging effect, conformational change, structural destability, molecular interactions to characterize the nature of the mutant protein^12^. Therefore, we provide a comprehensive understanding of the potentially deleterious SNPs that might be worth exploring for lung cancer pathogenesis.

## Materials and methods

### Lung cancer genes data retrieval

Genes responsible for three prominent types of lung cancers including Adenocarcinoma of lung (AC), Squamous cell carcinoma (SQCC) and small cell lung carcinoma (SCLC) were searched through previously published articles^5,13,14^. The present investigation did not involve human samples or participants, and the data generated are incorporated within this paper and are publically accessible.

### Protein-protein interaction networking

Proteins encoded by the selected genes were subjected to build PPI networks by using Cytoscape (v3.10.2) which is a Java based free software and can help in visualization of PPI networks with great ease^15^. Three different networks were built for three types of cancers for the selection of proteins with highest degrees. Most significant proteins contributing to different pathway aberrations which leads to lung cancers were identified.

### Exploration of SNPs

One of the most significant proteins identified during protein-protein networking, PTEN was searched in dbSNP database (https://ncbi.nlm.nih.gov/snp/) which is part of National Center for Biotechnology Information (NCBI). We filtered the initial search result with different filters (likely pathogenic, pathogenic, missense, somatic). Further, we retrieved information related to selected SNPs such as SNP ID, location, amino acid change. The protein sequence for this gene of interest was retrieved from UniProt database (https://www.uniprot.org/) (UniProt ID: P60484)^16^

### Identification of deleterious SNPs

We utilized online based seven different bioinformatics tools including SIFT (Sorting Intolerant From Tolerant) (https://sift.bii.a-star.edu.sg/www/SIFT_dbSNP.html) ^17–19^, SNPS&GO (https://snps-and-go.biocomp.unibo.it/snps-and-go/index.html) ^20^, PMut (https://mmb.irbbarcelona.org/PMut/analyses/new/) ^21^, PHD-SNP (https://snps.biofold.org/phd-snp/phd-snp.html) ^22,23^, PANTHER (https://www.pantherdb.org/tools/csnpScoreForm.jsp) ^24–26^, PROVEAN (http://provean.jcvi.org/seq_submit.php) ^27^, FATHMM (http://fathmm.biocompute.org.uk/cancer.html) ^28^. These programs predicted the effects of SNPs on functionality of PTEN protein. Merging results from all these databases ensured the accuracy of the results and helped to identify the potential SNPs as high risk to cause cancer. SIFT is a sequence homology-based method to forecast if a substitution would affect a protein’s phenotype at a certain location ^17,18^. SNPS&GO is a technique that uses the functional annotation of proteins to predict their relation with disease, based on their sequence. This is an SVM-based method that provides a GO-based score ^20^. PMut, a web-based tool can handle different steps such as feature computation, classifier selection, validation, and outcome analysis. These are based on python libraries. After evaluation using machine learning and other approaches, the web application provides a prediction score ranging from 0 to 1 ^21^. PhD-SNP uses SVM-based method using sequence information, profile information or hybrid method. SVM-Profile uses the output from the BLAST program. In PhD-SNP, the reliability index, or RI value, is calculated using the support vector machine’s output^22,23^. PANTHER calculates preservation during evolution. When a particular site of a protein is conserved, then there is highly chance of deleterious effect if it suddenly changed. This feature is taken into account when PANTHER- PSEP works ^26^.

PROVEAN (Protein Variation Effect Analyzer) is a tool that can forecast about the functional impact of protein variants. This tool computes delta score, assign a PROVEAN score and make prediction without any biasness^27^. FATHMM approach presents an unweighted/species-independent mechanism. This mechanism employs an iterative search process to automatically gather and align homologous sequences ^28^.

### Structural impact prediction of nsSNPs on PTEN

Seven different structural impact prediction and visualization tools were employed. These are MuPro (https://mupro.proteomics.ics.uci.edu/) ^29^, NetSurfP-2.0 ( https://services.healthtech.dtu.dk/services/NetSurfP-2.0/) ^30^, The mutation3D (http://www.mutation3d.org/) ^31^, Hope server (https://www3.cmbi.umcn.nl/hope/) ^32^, I-Mutant 2.0 (https://folding.biofold.org/cgi-bin/i-mutant2.0.cgi) ^33^, Missense3D (http://missense3d.bc.ic.ac.uk/missense3d/)^34^ , CASTp server (http://sts.bioe.uic.edu/castp/calculation.html) ^35^, DynaMut (https://biosig.lab.uq.edu.au/dynamut/) ^36^. These tools were utilized to identify structural impact and visual assessment of nsSNPs on PTEN protein structure. MuPro uses a different machine-learning approach from other tools which were used in the identification of deleterious SNPs. However, it is mainly based on support-vector machines that takes into account both structure and sequence-based information ^29^. NetSurfP-2.0^30^was used to predict the PTEN protein surface accessibility, secondary structure and disorders present in different locations. With mutation3D^31^ ,the 3D structure of the PTEN protein and the positions of mutations were visualized. Further, Mutation3D can be applied to seek whether a mutation is contained within a domain. HOPE (Have (y)Our Protein Explained) server ^32^ is a web-based service for analysis of mutations present on genes. I-Mutant2.0 has been fine-tuned to anticipate changes in protein stability resulting from mutations. It is a web server based on Support Vector Machines that is able to identify changes in protein stability caused by single-site mutations ^33^. The Missense 3D server is a freely available tool for structural investigation of missense variations^34^. The server can assess the effect of mutation on the wild protein. The pdb files of refined 3D models were uploaded to CASTp (Computed Atlas of Surface Topography of proteins) server ^35^. The server calculates the cavities, channels as well as the pockets. DynaMut is one of the widely known comprehensive tool for understanding protein motion and flexibility ^36^. To assess how point mutations affect protein stability, it combines normal mode parameters with optimized graph-based signatures.

### Conservation profile analysis of PTEN

To locate the evolutionary conserved sites in the amino acid sequence of PTEN gene, the Consurf web server (http://consurf.tau.ac.il) was used ^37^. An amino acid’s degree of evolutionary conservation in a protein or a nucleic acid’s degree of conservation in DNA/RNA shows a trade-off between the macromolecule’s overall need to maintain its structural integrity and function and its inherent tendency to mutate. This tool automatically finds homologues based on a query sequence or structure, deduces their multiple sequence alignment, and builds a phylogenetic tree that shows their evolutionary relationships.

### Determining the locations and features of nsSNPs on PTEN domians

To pinpoint the locations of nsSNPs on different domains of PTEN, we used InterPro and PROSITE. The InterPro database^38^ (https://www.ebi.ac.uk/interpro/) classifies the families of protein sequences and finds functionally significant domains and conserved regions. PROSITE (https://prosite.expasy.org/) is a collection of protein families and domains ^39^. It is predicated on the finding that, despite the vast array of diverse proteins, the majority of them can be classified into a small number of families according to sequence similarities.

### Three-dimensional (3D) modeling of native and mutant PTEN protein

To build 3D model of native and mutant PTEN protein, we utilized three tools including HHPred, AlphaFold Server and trRosetta. The protein models were built using the protein FASTA sequences as input. HHpred (https://toolkit.tuebingen.mpg.de/tools/hhpred) is a fast server that generates multiple query alignments with selected protein templates chosen from the search results, pairwise query-template alignments, and MODELLER software-calculated 3D structural models ^40^. AlphaFold□3 (AF3) (https://alphafoldserver.com/) predicts the structure with improved protein structure accuracy and understanding the structure of protein-protein interactions ^41^. The trRosetta server (transform-restrained Rosetta) (https://yanglab.qd.sdu.edu.cn/trRosetta/) is a web-based platform that uses deep learning and Rosetta to predict protein structures quickly and accurately^42^.

### Model verification and refinement

PROCHECK (https://saves.mbi.ucla.edu/) was used to verify the constructed structures using three distinct methods in order to evaluate their quality^43^. This tool assesses the structural integrity of a protein by checking the geometry of residues. The Ramachandran plot was used to check the model structure’s quality, and the SAVES web server was used to certify the structure’s further verification. If 90% of the residues of a protein structure are located in the preferred region within the Ramachandran Plot, then the structure is considered good ^44^. These methods are essential to comprehending the three-dimensional protein models. The GalaxyRefine server (https://galaxy.seoklab.org/cgi-bin/submit.cgi?type=REFINE) was used to increase the accuracy of PTEN wild type and mutant 3D models of HHPred ^45^.

### Molecular dynamic simulation

GROMACS (GROningen MAchine for Chemical Simulations) simulation software (Version 2024.3) was used for molecular dynamics simulation of the wild type PTEN and mutant types for 100 ns ^46^. Different analyses including Hydrogen-hydrogen bonds (HHBonds), the radius of gyration (Rg), root mean square deviation (RMSD), root mean square fluctuation (RMSF), and solvent accessible surface area (SASA) were carried out using the modules of GROMACS package. The plots for the results of these analyses were generated using QtGrace software (v026).

### Ligand selection

To demonstrate the mutational impacts on interaction of the PTEN protein, we selected ligands that have the property of interacting with PTEN from DrugBank. One specialized site for pharmaceutical research is DrugBank ^47^ which provides essential information on how several proteins interact with various medicines. Numerous drug repurposing trials and thousands of experimental drug clinical trials constantly add new data ^48^.

### Molecular docking analysis

To determine the effect of deleterious point mutations over the binding affinity of PTEN, we performed molecular docking using the AutoDock Vina tool. Predicting the bound conformations and binding affinity is one of the main targets^49,50^. The PDB format of ligand and receptor proteins were converted into pdbqt format using AutodockVina. We visualized the position of SNP on mutant protein using Pymol software and set parameters targeted to the specific region (where the wild type residue has been changed into mutant type) for both mutant and wild proteins. The BIOVIA Discovery Studio^51^ tool was used to visualize the docking result.

## Results

### An intersection of lung cancer cell types found PTEN to be more prevalent

Mutations in genes that control different pathways namely RTK, RAS/RAF, PI3K/AKT, LKB1/AMPK, TP53, MYC, Developmental pathway are responsible for different types of lung cancer ^5^. Some of the genes may contribute to multiple lung cancer types. The genes were identified by searching through literature and listed in **Supplementary Table 1**.

PPI networks illustrate the interactions between various genes. For each form of cancer, the top 10 genes with the highest degree were displayed by Cytoscape. KRAS, TP53, PIK3CA, PTEN, BRAF, MYC, EGFR, NF1, ERBB2 and NRAS were among the top 10 genes predicted for adenocarcinoma of lungs (**Figure 1A and Supplementary Table 2**). On the other hand, Cytoscape visualized TP53, PTEN, PIK3CA, KRAS, EGFR, NOTCH1, AKT1, NF1, ERBB2, HRAS as top 10 genes responsible for squamous cell carcinoma (**Figure 1B and Supplementary Table 3**). Moreover, predicting top 10 genes for small cell lung carcinoma revealed PTEN, TP53, MYC, CREBBP, CCNE1, MYCN, RB1, EP300, KMT2A, FGFR1 with highest degree (**Figure 1C and Supplementary Table 4**). Genes ranked with the highest degree might function as hub genes that have a major impact on several pathways’ activities and further an intersection network analysis of the top ten genes from three forms of lung cancer illustrated that TP53 interacts with PTEN (**Figure 1E**). In addition, the Venn diagram demonstrated that only two genes, TP53 and PTEN, are present in three kinds of lung cancer (**Figure 1D**). PTEN was chosen for further downstream analysis on the effects of its mutations on various lung cancer types since it was one of the genes that scored first in the network analysis of all three types of lung cancer using the degree method (**Supplementary Table 2-4**).

**Figure 1:**
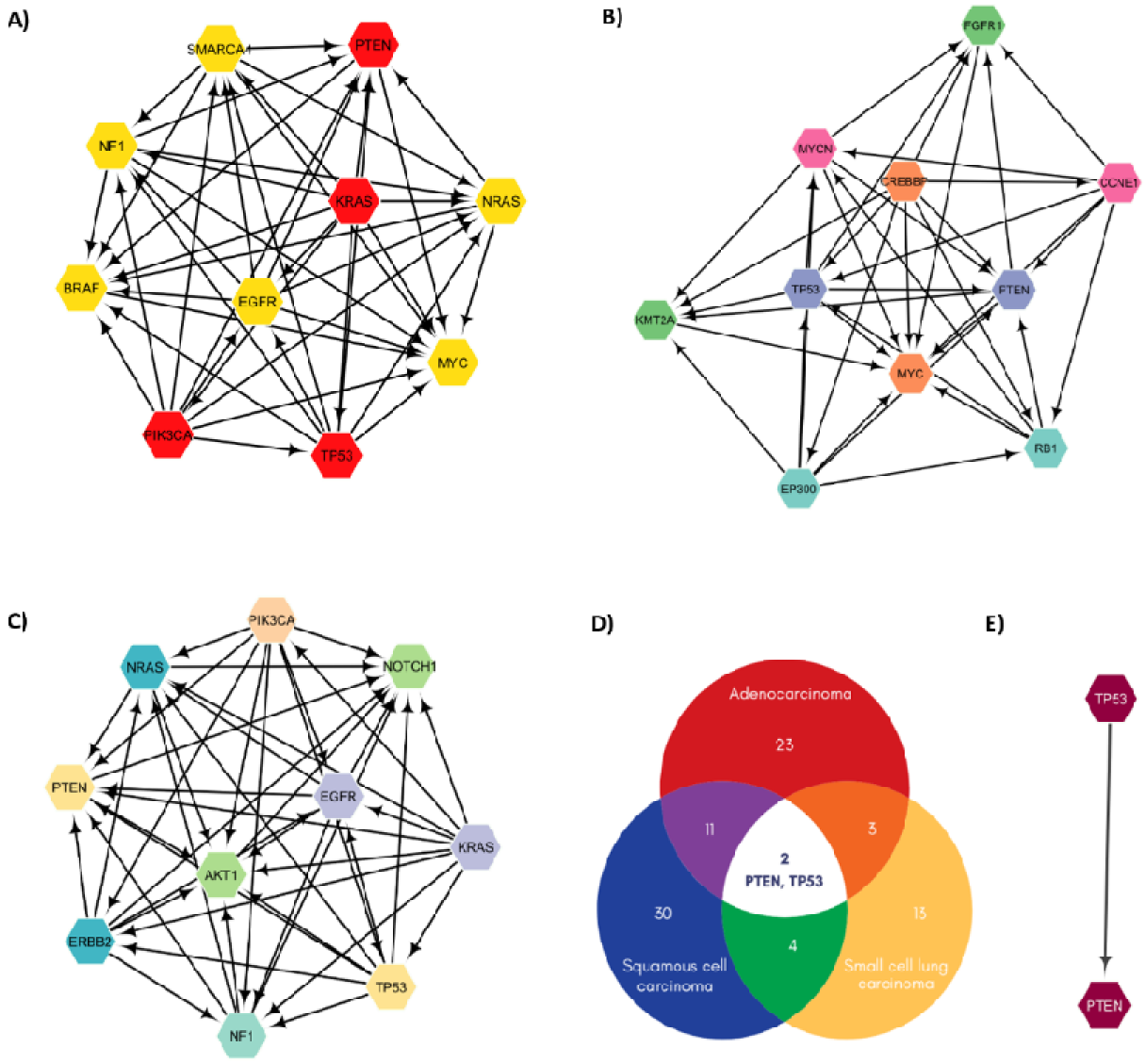
Highly connected genes identification of the different lung cancer cell types. Top 1 genes ranked by degree method responsible for A) Adenocarcinoma of lungs, B) Squamous cell carcinoma, C) Small cell lung carcinoma. The arrows from nodes to nodes represent the interaction of one gene with another. D) Representation by Venn diagram to pinpoint most significant genes in three types of lung cancer. E) Intersectio network analysis of all top genes.

### Computational assessment revealed 15 nSNPs as deleterious

PTEN was looked up in the National Center for Biotechnology Information’s (NCBI) dbSNP database. We filtered 163 SNPs out of 43855 SNPs found through initial search result with different filters (likely pathogenic, pathogenic, missense, somatic).

Seven tools were utilized including SIFT, SNPs&GO, PMut, PhD-SNP, PANTHER, PROVEAN and FATHMM to determine the functional implications of nsSNPs on the PTEN gene product **(Supplementary Table 5-11)**. Out of total 163 nsSNPs, only 15 were commonly identified as deleterious by all tools which are presented in **Table 1 and Figure 2**. These tools labeled the nsSNPs as deleterious, disease, probably damaging, decrease stability or cancer. These 15 noteworthy nsSNPs were taken into account for the subsequent filtering step.

**Figure 2:**
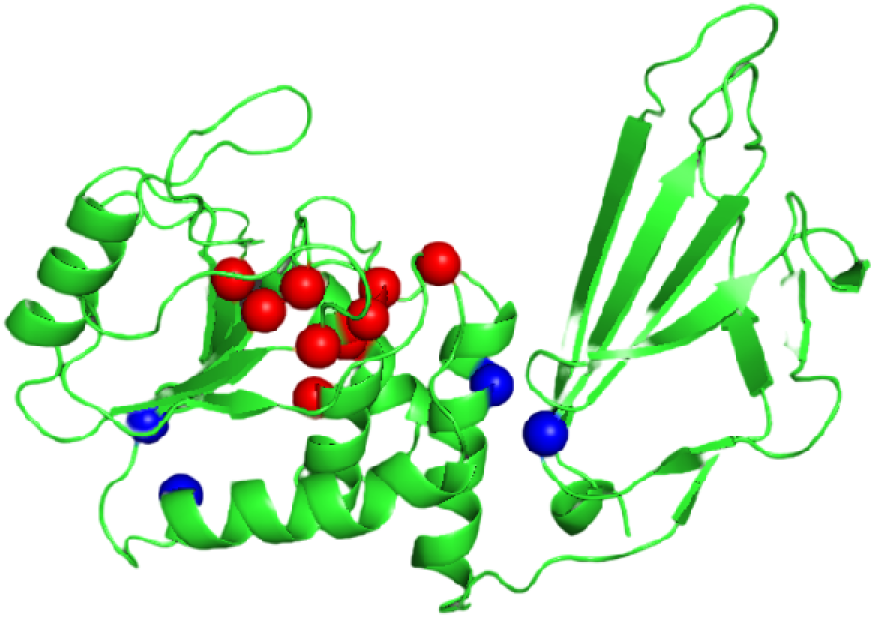
Visualization of the 15 mutations in the protein structure. Mutation 3D tool used the PDB model 1D5R (AA range: 14-281) which showed the presence of selected mutations. Red balls indicate the locations of G129E, H123R, C124R, R130G, M35R, L70P, R130Q, H93R, G132V, I135T mutations and blue balls indicate the locations of R173C, R173H, V119L, L112P, D252G mutations.

**Table 1:**
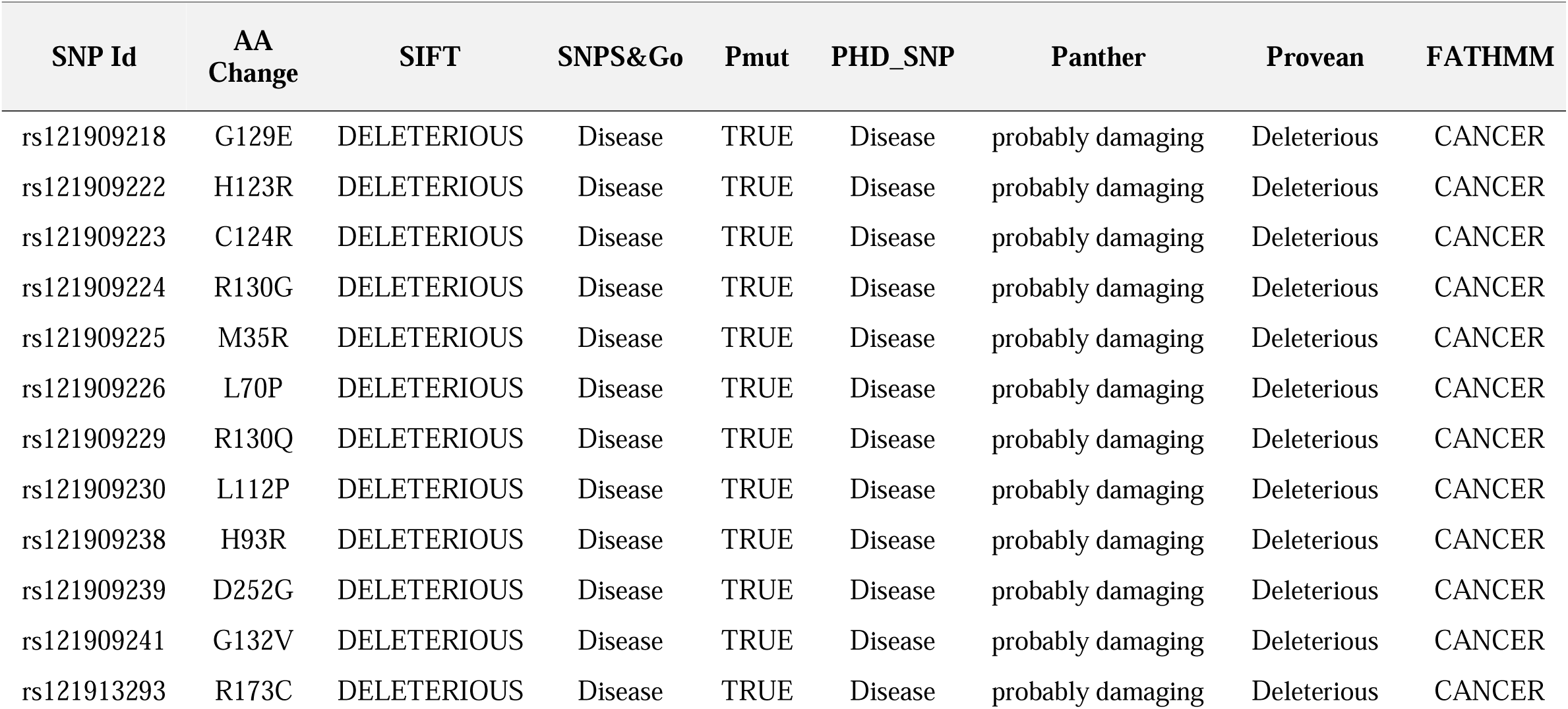
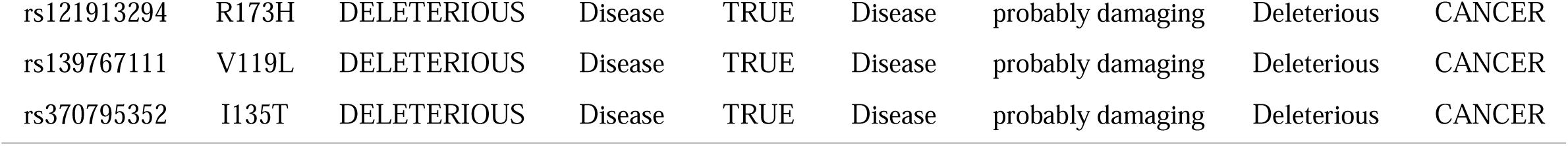
Identification of deleterious nsSNPs using different types of algorithms.

### Computational assessment revealed seven mutations could lead the conformational shift

Fifteen nsSNPs projected to be harmful by seven different tools underwent structural impact analysis.

NetsurfP-2.0 was used to evaluate the stability and accessibility of native and mutant proteins for each of the 15 variations. The percentage prediction of NetSurfP-2.0 indicates the buried or exposed status of a residue inside the protein structure (**Table 2**). Two nsSNPs (R173C, R173H) were discovered to be exposed in the wild type (**Supplementary Figure 1**), while the remaining nsSNPs were buried. R173C was buried in the mutant, but R173H remained exposed in it a well. Two more nsSNPs (L112P, H93R) were also exposed in the mutant (**Supplementary Figure 2**), while the other nsSNPs maintained buried status.

**Table 2:**
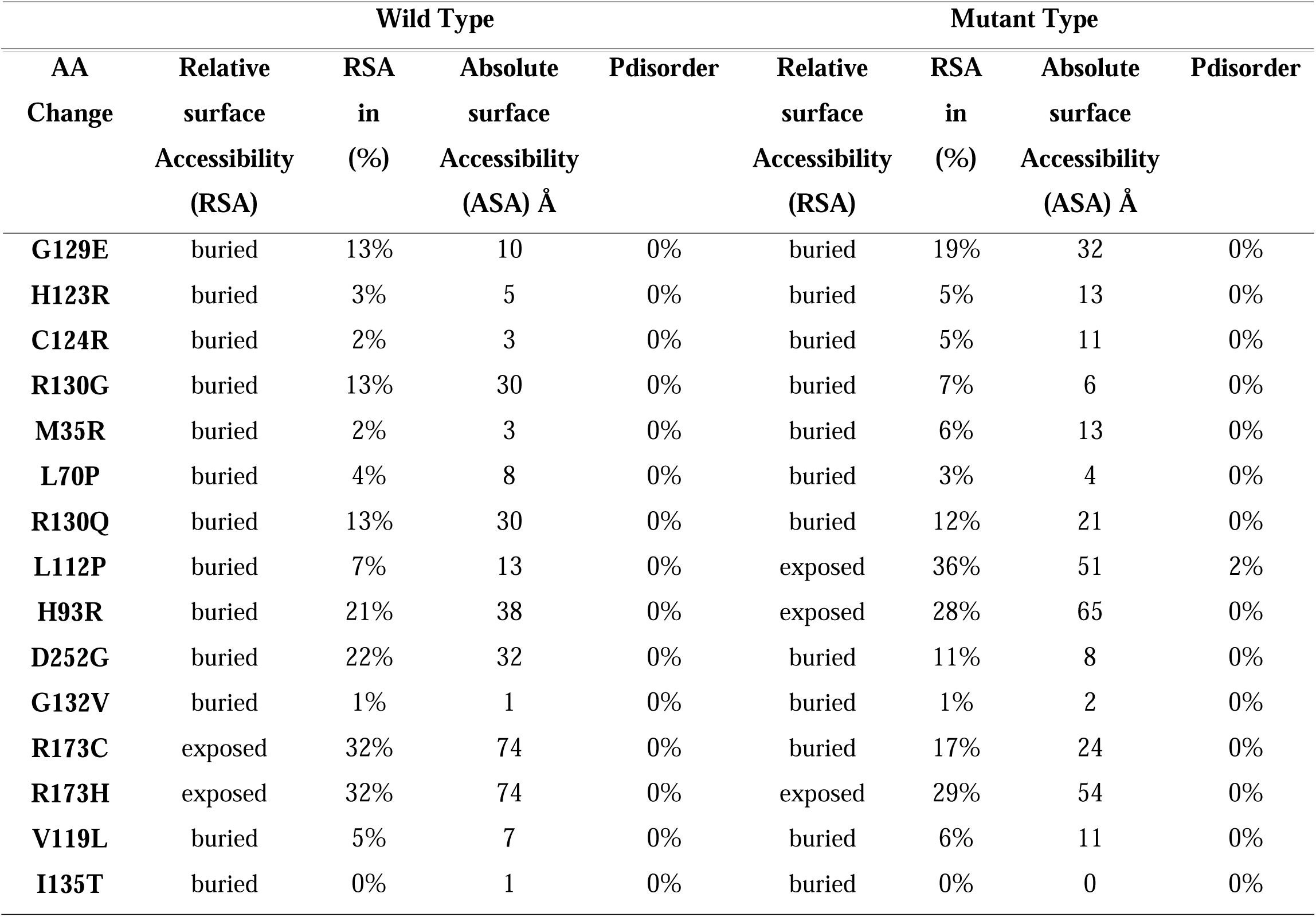
Surface accessibility predicted by NetsurfP-2.0 for native and mutant of PTEN.

The results of I-Mutant 2.0 projected the changes in PTEN stability in terms of RI and free energy change values (DDG), which involved introducing point mutations in the PTEN protein. The results found that 14 of the 15 harmful nsSNPs reduced stability (**Supplementary Table 12**). Another tool, MuPro was also utilized to forecast the effects of single-site amino acid mutation on the stability of proteins. All of the 15 harmful nsSNPs were predicted to have reduced stability (**Supplementary Table 13**). On the other hand, Missense 3D tool predicted 9 SNPs (H123R, C124R, R130G, M35R, L70P, R130Q, H93R, D252G, G132V) to be damaging while the rest to be neutral (**Supplementary Table 14**) Additionally, investigation with HOPE server resulted with 7 mutant residues (G129E, H123R, C124R, M35R, H93R, G132V, V119L) larger than the wild type, while 8 mutant residues were smaller than the wild type (R130G, L70P, R130Q, L112P, D252G, R173C, R173H, I135T). The effect on stability is listed on the **Supplementary Table 15**. CASTp predicted pockets’ area and volume for mutant and native proteins. The biggest pockets spanned across the important domain regions which varied among different mutants and native PTEN protein indicating possible conformational shift and functional impact (**Figure 3 and Supplementary Table 16**).

**Figure 3:**
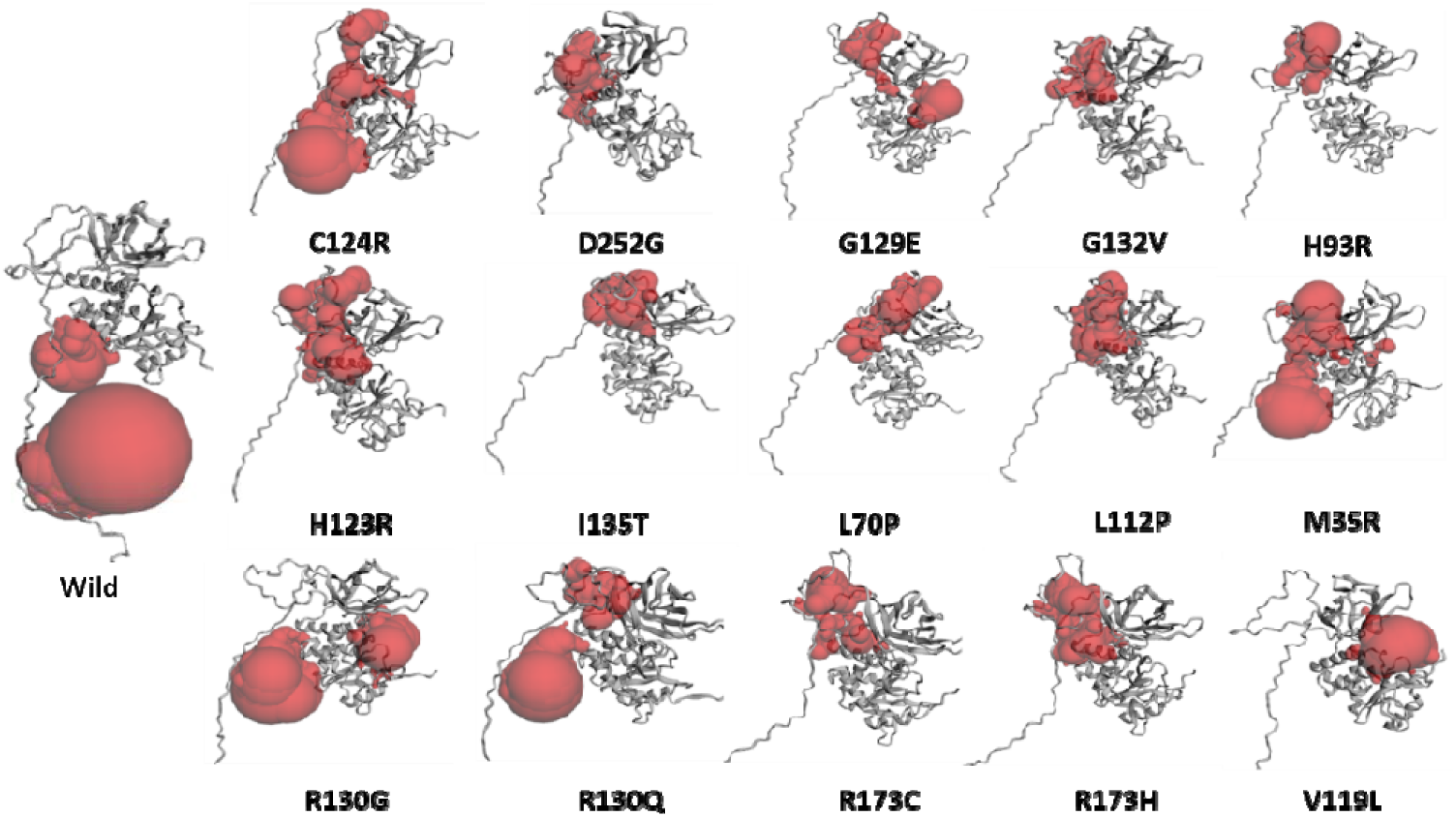
Active pocket of wild and mutant structure. CASTp predicted biggest pockets for native and mutant proteins spanning the important domain region are visualized. Red color indicates the presence of pockets. Significant difference in pockets location and sizes of mutants from wild type is identified.

The DynaMut server provided the analysis report of increase or decrease in stability (ΔΔG) of nsSNPs. These were expressed in kcal/mol and a destabilizing influence is known by negative value, and a stabilizing effect is known by a positive value. The more destabilizing the mutation is for the protein, the lower the ΔΔG value. Ten nsSNPs (C124R, D252G, H123R, I135T, L70P, L112P, M35R, R130G, R173C, R173H) showed destabilizing impact on the protein structure. Five nsSNPs (G129E, G132V, H93R, R130Q, V119L) had shown stabilizing impact on protein structure (**Supplementary Table 17**). The results from these tools identified seven high-risk mutations out of fifteen with most conformational shift (**Table 3**).

**Table 3:** Prediction of high-risk PTEN nsSNPs according to conformational assessment.

| Mutation | I-Mutant 2.0 | Consurf | MuPro | HOPE server | DynaMut |
| --- | --- | --- | --- | --- | --- |
| **H123R** | Decrease | Conserved | DECREASE stability | Probably damaging | Destabilizing |
| **C124R** | Decrease | Conserved | DECREASE stability | Probably damaging | Destabilizing |
| **R130G** | Decrease | Conserved | DECREASE stability | Probably damaging | Destabilizing |
| **M35R** | Decrease | Conserved | DECREASE stability | Probably damaging | Destabilizing |
| **L70P** | Decrease | Conserved | DECREASE stability | Probably damaging | Destabilizing |
| **D252G** | Decrease | Conserved | DECREASE stability | Probably damaging | Destabilizing |
| **R173C** | Decrease | Conserved | DECREASE stability | Probably damaging | Destabilizing |

### nsSNPs of PTEN revealed their location in functional domains

InterPro, a tool for identification of protein domains, active sites and other features can do the job of prediction by utilizing 13 protein signature databases. It predicted six domains which were summarized (**Supplementary Table 18**) and visualized (**Supplementary Figure 3**). Additionally, PROSITE predicted domains over the full range of protein (**Supplementary Figure 4**).

### nsSNPs of PTEN are found in conserved regions

The evolutionary conservation of the amino acid residues in the PTEN protein was computed using the ConSurf online tool in order to investigate the possible effects of the 15 nsSNPs that were shortlisted in upstream steps. The findings were displayed as a protein sequence visualization ranging 1-403 amino acids, emphasizing the conservation status by showing different colors.

ConSurf output revealed that 12 out of the 15 nsSNPs had a conservation score of 9, indicating that they were highly conserved residues. Only three nsSNPs were expected to be have conservation score of 8. Variants located in these conserved regions are considered to be highly damaging to the protein, as referred to as those located in non-conserved sites. The deleterious predictions for every residue including each selected SNP by ConSurf are summarized in **Figure 4**.

**Figure 4:**
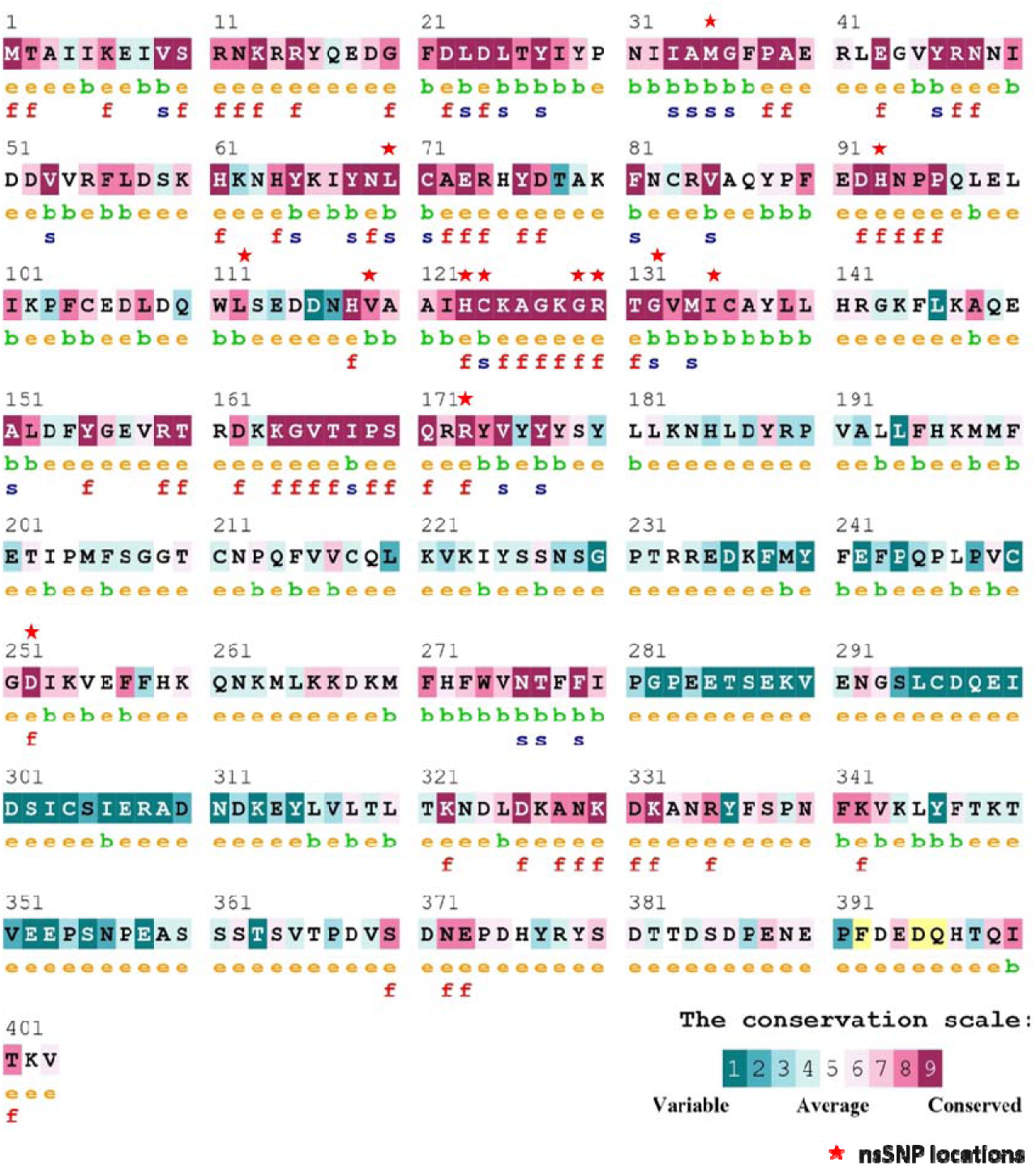
Evolutionary conservation profile analysis of PTEN by Consurf web. nsSNP locations are indicated by red star marks. Here, ‘e’ = an exposed residue, ‘b’ = a buried residue, ‘f’ = a predicted functional residue (highly conserved and exposed), ‘s’ = a predicted structural residue (highly conserved an buried), ‘x’ = insufficient data where calculation for that site was performed on less than 10% of the sequence.

### Structural impact of seven mutations predicted the destable nature of PTEN

Using the HHPred server, AlphaFold, trRosetta server, the protein 3D model was built in PDB format. HHPred server usage required a MODELLER key which was utilized for generating the models.

The quality of 3D models was validated by inspecting the Ramachandran plot generated by th PROCHECK & ERRAT server from SAVES. The findings indicated that over 90% of the residues of the homology model belonged to the most favored regions for both the wild and mutant version. Although ERRAT score predicted to define overall quality score of different modeling servers were fluctuating, none of them were appropriate in 3D model structure. PROCHECK validated this assessment by numerical values which showed the average percentage of residues in most favored region for HHPred was 88% (86-89% range), for Alphafold was 82.74% (81-84% range), for trRosetta was 86.57% (85-89% range). The values provided by ERRAT and PROCHECK server for HHPred, Alphafold and trRosetta predicted models are listed in **Supplementary Table 19-21**. By taking this information into account and to remove complexity, only pdb files from HHpred were utilized to refine them by GalaxyRefine server.

First model out of five models were chosen for every entry. The chosen models’ different predicted metrics are listed on **Supplementary Table 22**. After refinement, the models were inspected with SAVES server’s PROCHECK and ERRAT again and showed significant improvement in the average percentage of residues in most favored region. Twelve models showed the average percentage of residues in most favored region over 90% after refinement, indicating appropriate 3D model (**Supplementary Table 23**). Four models (L70P, L112P, R173H, V119L) showed the average percentage of residues in most favored region nearly but below 90% after refinement, indicating inappropriate 3D model. However, these percentages were still above the percentages predicted before refinement **(Supplementary Figure 5)**. As a result, all of these models were utilized for further steps. Additionally, all models received an ERRAT score greater than 50, indicating better quality for all models predicted by the GalaxyRefine server.

### Molecular dynamic simulation confirmed the destabilty of the PTEN structure

We performed MD simulation to evaluate the overall stability of PTEN and the variants using different metrices such as HHBond, Rg, RMSD, RMSF and SASA. For understanding the protein stability, folding and interactions, HHBond analysis is performed. Higher hydrogen bond numbers are indicators of more stable structures. During the experiment, C124R, L70P and H123R showed higher hydrogen bonds compared to wild type. On the contrary, R130G showed lower hydrogen bonds compared to wild type. Additionally, M35R, D252G and R173C showed almost similar hydrogen bonds when compared to wild type (**Figure 5**).

**Figure 5:**
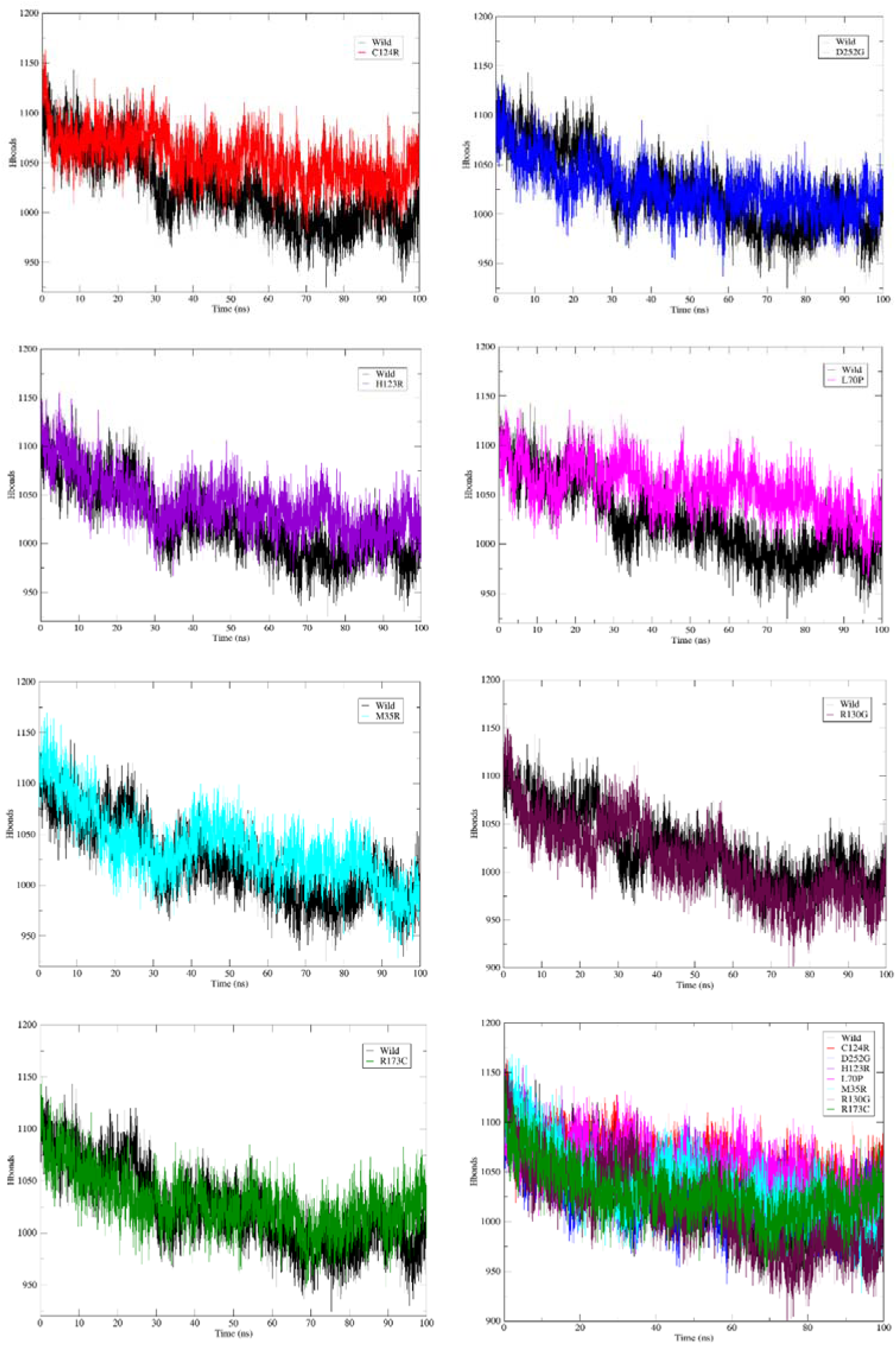
HHBond analysis of PTEN wild type and mutant structure. Wild type (Black line), C124R (Red line), D252G (Blue line), H123R (Violet line), L70P (Magenta line), M35R (Cyan line), R130G (Maroon line), R173C (Green line).

The radius of gyration value corresponds to the compactness of the protein. This analysis was performed to know if the PTEN protein and its mutant types were correctly folded or not. Out of all mutants under experiment, C124R, H123R and L70P showed more Rg value than the wild type over 100 ns time indicating their significant structural changes or denaturation due to the presence of SNP (**Figure 6**). Root Mean Square Deviation (RMSD) analysis was performed to analyze PTEN protein and its mutants’ deviation in state from their initial states. In the experiment, C124R and H123R showed maximum RMSD profile over the time compared to the wild type. On the other hand, M35R, D252G, R173C possessed higher RMSD values than wild type in most of the time with periodic fluctuations indicating slight deviations. In contrast, L70P and R130G displayed significant fluctuations in RMSD value although R130G almost followed the trajectory of wild type over 100 ns time (**Figure 7**). Root Mean Square Fluctuation (RMSF) analysis was carried out to understand the flexible regions of native PTEN and the selected mutants. All of the mutant proteins showed higher flexibility in their respective mutant regions which might be an effect of the presence on mutant on the overall structure of the protein (**Figure 8**). Solvent Accessible Surface Area (SASA) analysis was done to know the surface area of PTEN and its variants which is accessible to external solvents. Higher SASA value is linked with the destabilization of the protein structure due to solvent accessibility. During the MD simulation, C124R, H123R and L70P showed higher SASA value than the wild type. Meanwhile, D252G had almost similar SASA value compared to wild type. However, M35R and R130G had slightly lower SASA values than the wild type in the end (**Figure 9**)

**Figure 6:**
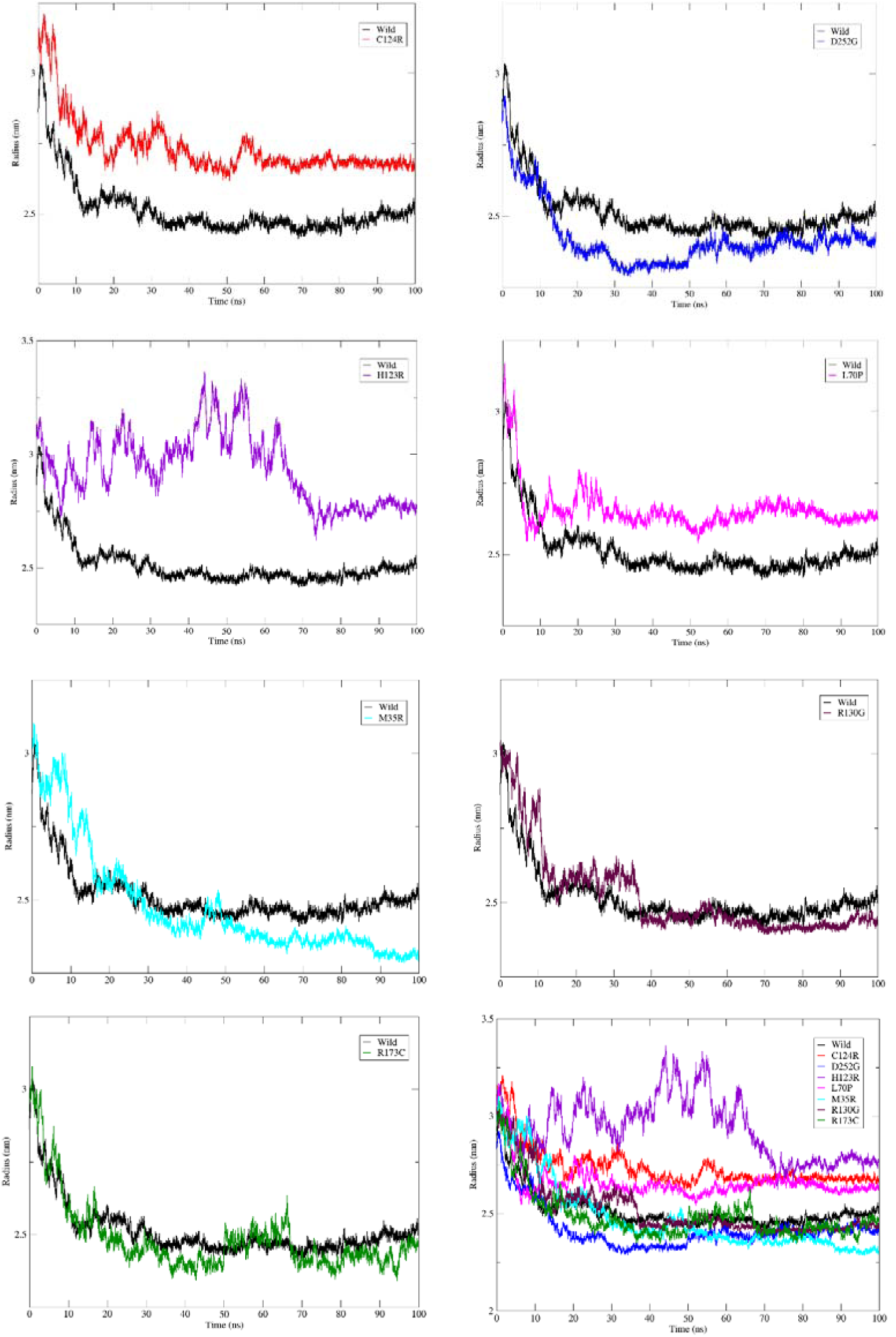
The radius of gyration (Rg) analysis of PTEN and mutant structure. Wild type (Black line), C124R (Red line), D252G (Blue line), H123R (Violet line), L70P (Magenta line), M35R (Cyan line), R130G (Maroon line), R173C (Green line).

**Figure 7:**
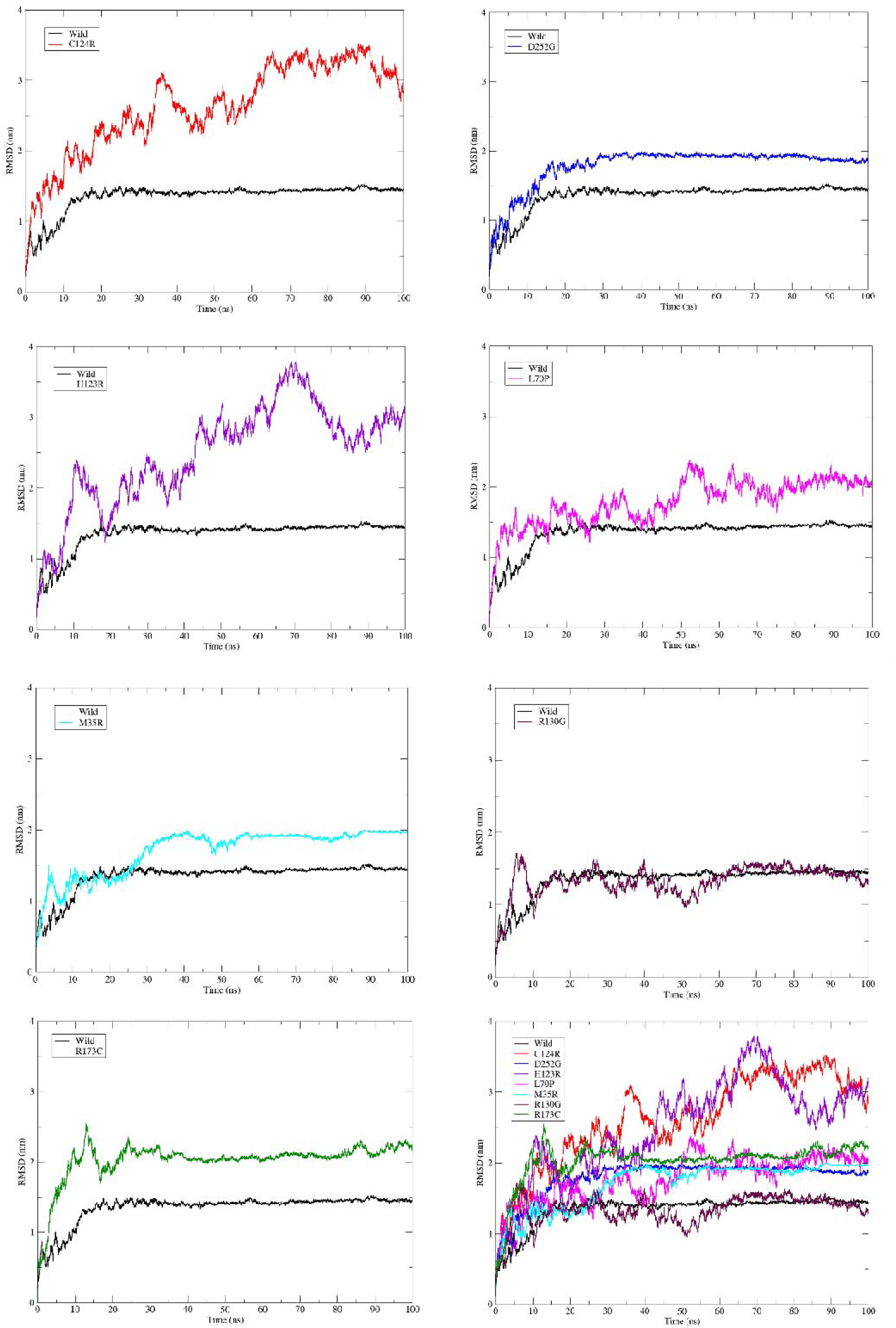
RMSD analysis of PTEN wild type and mutant structure. Wild type (Black line), C124R (Red line), D252G (Blue line), H123R (Violet line), L70P (Magenta line), M35R (Cyan line), R130G (Maroon line), R173C (Green line).

**Figure 8:**
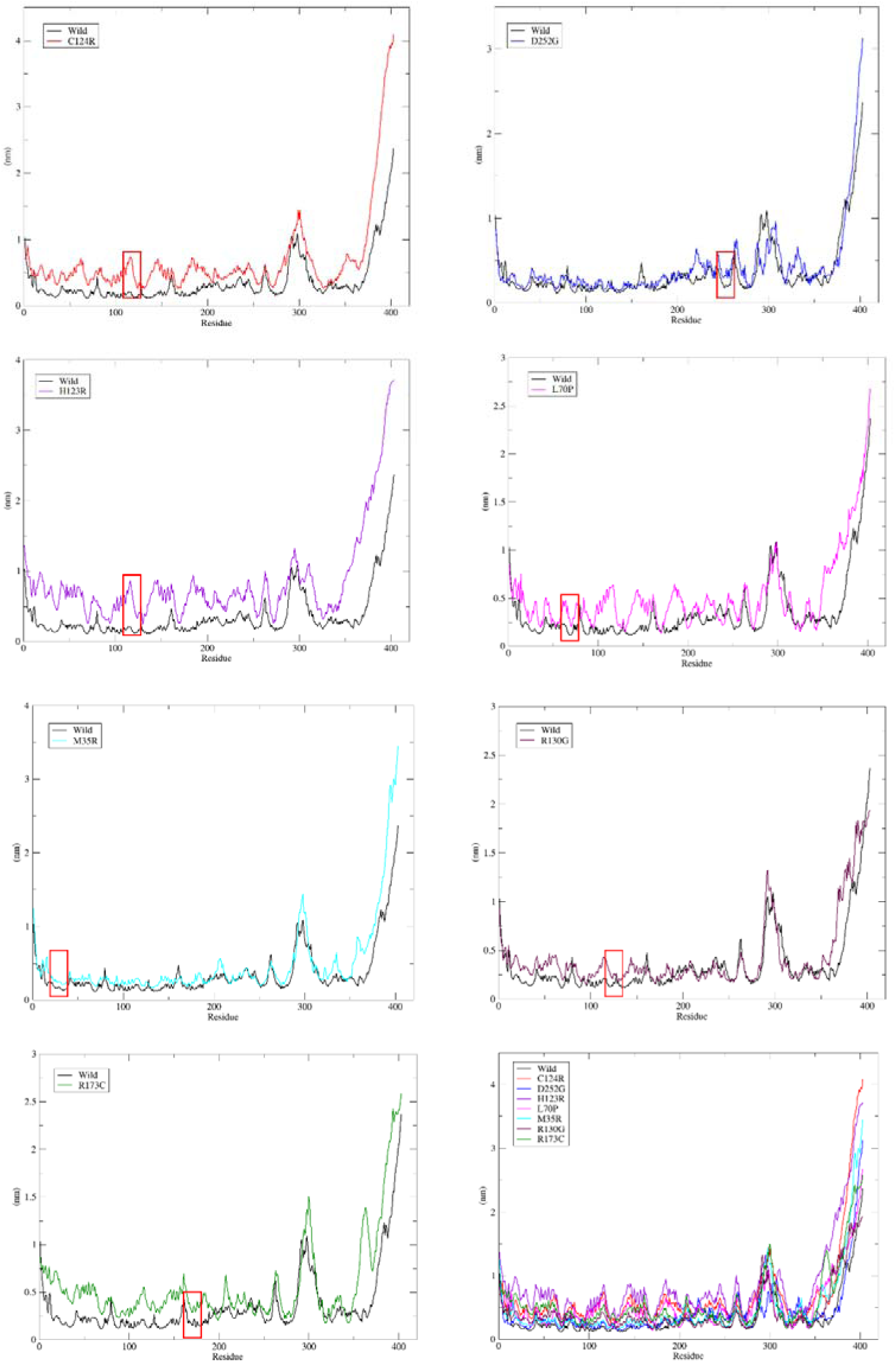
RMSF analysis PTEN of wild type and mutant structure. Wild type (Black line), C124R (Red line), D252G (Blue line), H123R (Violet line), L70P (Magenta line), M35R (Cyan line), R130G (Maroon line), R173C (Green line). Approximate mutant positions are showed in red box.

**Figure 9:**
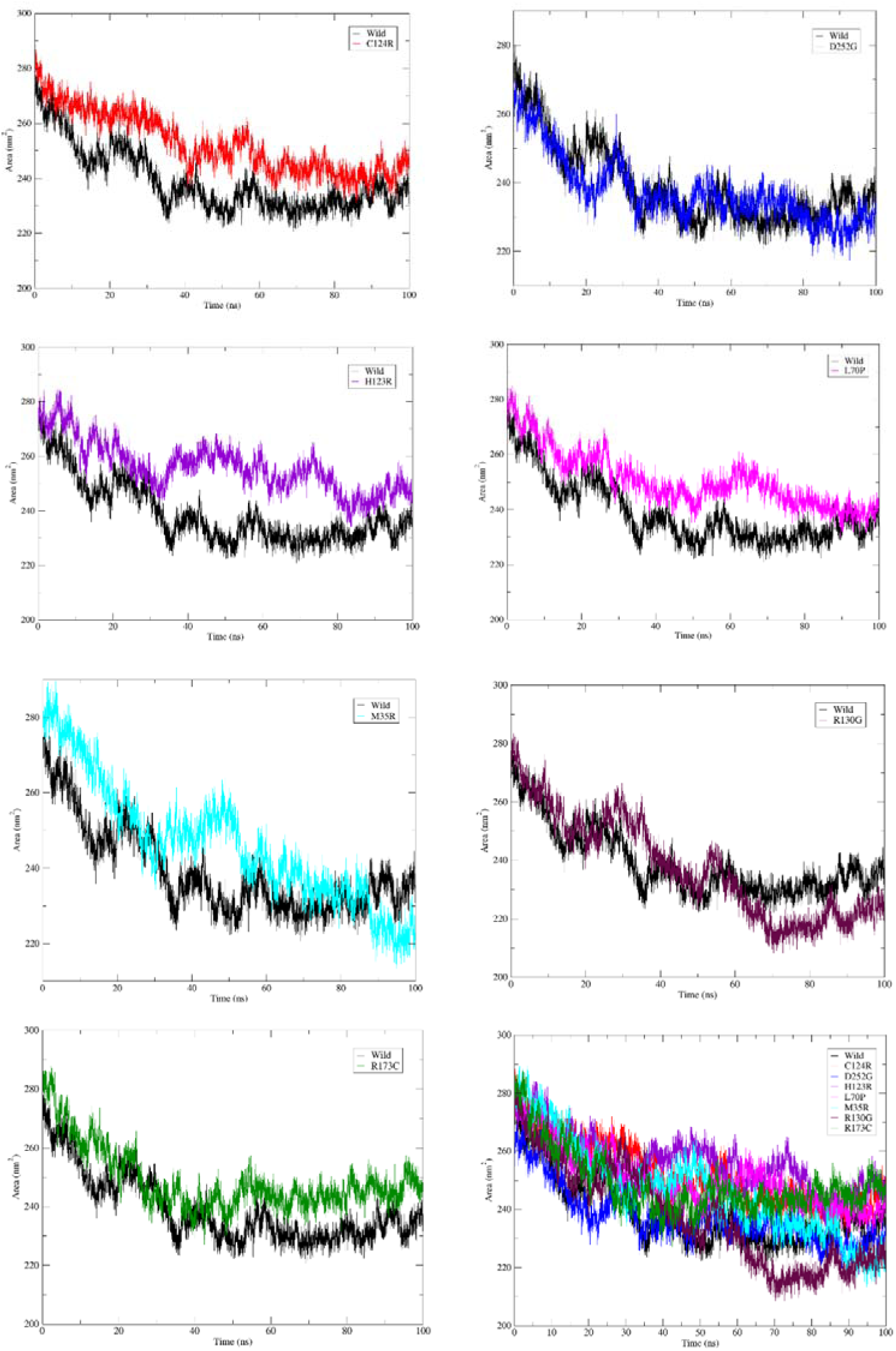
SASA analysis of PTEN wild type and mutant structure. Wild type (Black line), C124R (Red line), D252G (Blue line), H123R (Violet line), L70P (Magenta line), M35R (Cyan line), R130G (Maroon line), R173C (Green line).

### Drug binding experiments validated the confirmational change of the structure

The drug binding analysis confirmed the effect of deleterious point mutations over the binding affinity of PTEN. The ligand selected was Fostamatinib which acts as inhibitor of PTEN. The same gridbox parameters selected for both wild and mutant model (**Supplementary Table 24**) were utilized in docking analysis and displayed varied binding affinity for both wild type and mutants. The wild type of H123R and R173C showed significant results with mostly conventional hydrogen bond and alkyl interactions in the selected region while the mutant model displayed highest variation in RMSD compared with the wild type (**Supplementary Table 25**). Additionally, the interactions patterns were changed due to the generation of bonds with different residues in the same region **(Figure 10)**.

**Figure 10:**
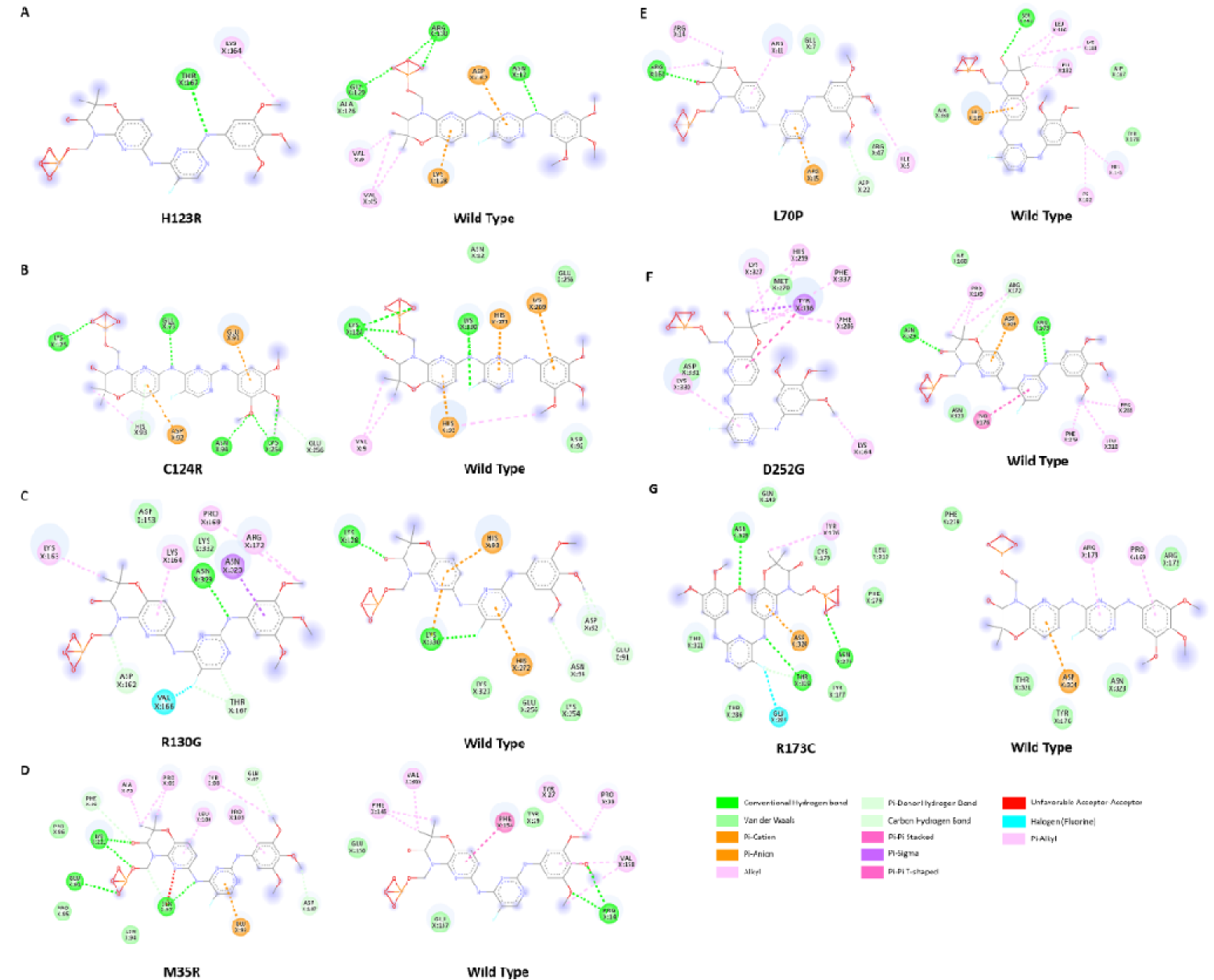
Visualization of amino acid interactions in same binding region of Fostamatinib with the amino acids of PTEN mutant and wild types, A) H123R, B) C124R, C) R130G, D) M35R, E) L70P, F) D252G, G) R173C.

## Discussion

Herein, we detail the extensive collection of PTEN nsSNPs and offer insights into the structural impact of genetic variants by functional and structural analysis. From a pool of around 43,855 nsSNPs, we were able to isolate 15 mutations that could provide insight into the genetic process behind the structural effect. In addition, molecular simulation and drug binding later supported the confirmational modification, and seven mutations ultimately confirmed the structure’s destability.

Proteins that function under human physiological conditions must maintain their structural integrity and functionality^52^ .When a protein exhibits structural instability, its associated functions are consequently compromised and lead to disease such as cancer **(Supplementary Table 1).** Mutations in P53 are generally recognized as indicative of lung cancer prognosis, whereas the causal role of PTEN in lung cancer is potentially significant; nonetheless, research on mutations in this gene remains insufficiently examined. Our findings further emphasizes that PTEN is a highly interconnected gene among NSCLC, SCLC, and adenocarcinoma **(Figure 1 and Supplementary Table 2-4).** The selection of the PTEN gene is corroborated by multiple studies, where similar methodology was employed to identify the other therapeutic targets ^53–56^. Having found 15 detrimental nsSNPs of PTEN (**Table 1**, **Figure 2 and Supplementary Table 5-11**), we employed a different array of bioinformatics techniques **(Supplementary Table 12- 17)** that predicted several parameters such as stability for function, secondary structure, and surface accessibility, which critically evaluated the functional divergence of the mutant PTEN proteins from the native variant (**Table 2 and Supplementary Figure 1,2**). Furthermore, conservation analysis demonstrated the potential for deleterious mutations in the conserved areas of PTEN, suggesting the harmful nature of the mutations in protein’s function (**Figure 4**). These insights have been proven as all the identified nsSNPs reside inside functional domains that may result in loss of function due to the specific mutation (**Supplementary Figure 3, 4 and Supplementary Table 18**).

However, the structural impact is crucial to make such claims; therefore, the presence of the mutations in the mutant structures underwent verification and refinement (**Supplementary Table 19-23**) to produce a configuration that closely resembles the realistic scenario of the wild type structure (**Supplementary Figure 5**). The results suggest the different binding locations of the structure if the mutation is present compare to the wild type structure. Analysis of interatomic interactions elucidated the diminishing stability following the introduction of mutations (**Supplementary Table 17**). The emergence of novel pockets on the mutant proteins, in contrast to the natural type, indicated visible structural modifications and altered functional activity (**Figure 3 and Supplementary Table 18**).

Subsequent molecular dynamics of both wild-type and selected mutant (**Table 3**) structures indicated the destabilising impact of the mutations, as evidenced by the 100 ns molecular dynamic simulation. The impacts are assessed under *in silico* physiological conditions, and the results of these analyses are anticipated to be reproducible in wet lab environments. The structural stability is diminished due to the mutation’s presence compared to the wildtype structure, as anticipated by the variants utilising several metrics such as HHBond, Rg, RMSD, RMSF, and SASA. The compactness, deviation, fluctuation, solvent accessibility, and hydrogen bond interactions of the mutant structure exhibited a distinct structural folding pattern compared to the wild-type structure, as confirmed by the time-dependent analysis over 100 ns (**Figure 5-9**). This indicates that the atoms and molecules possess different organisational orientations due to the mutation’s presence. Consequently, binding studies with Fostamatinib **(Supplementary Table 24)**, a specific inhibitor of PTEN, revealed four deleterious nsSNPs (H123R, M35R, D252G, R173C) exhibiting significantly lower binding affinity relative to the wild type. The other three mutations (C124R, R130G, L70P) showed the moderate difference in binding affinity with wild type (**Supplementary Table 25**). Furthermore, the amino acid residues exhibited a completely altered pattern compared to the wild type structure. All mutations exhibit distinct interactions of amino acid residues compared to the wild-type structure in the same binding site. Hence, these mutations confirmed almost or complete altered interaction pattern than the wild type with Fostamatinib indicating major conformational change in structure (**Figure 10**).

All cross-referenced data corroborated the destabilisation of the structure in the presence of mutations, thereby may facilitate the impairment of the PTEN function in relation to the anomaly in inducing functionality. In order to better understand the function of PTEN nsSNPs in lung cancer, our study distinguished these 7 nsSNPs from other mutations found in the database. Through a variety of molecular approaches, it has also made it easier to identify deleterious variations. In addition, the data can be used to personalized treatments for PTEN gene mutations.

## Conclusion

This study discovered seven mutations in the PTEN gene that may disrupt protein function. The present study notably simulated the mutation inside the PTEN structure, confirming the conformational shift and then validating the alteration in folding patterns. Consequently, these identified mutations significantly impact lung cancer prognosis. Ultimately, this study facilitates additional exploration of the biological effects and mechanisms of these nsSNPs.

## Data Availability

All data generated or analyzed during this study are included in this published article.

## Authorship contribution statement

**Mohammad Uzzal Hossain:** Data curation, Formal analysis, Conceptualization, Methodology, Software, Visualization, Writing – original draft. **Mohammad Nazmus Sakib:** Data curation, Formal analysis, Conceptualization, Methodology, Software, Visualization, Writing – original draft. **A.B.Z. Naimur Rahman:** Data curation, Formal analysis, Methodology, Software, Visualization, Writing – original draft. **SM Sajid Hasan:** Data curation, Formal analysis, Methodology, Software, Visualization, Writing – original draft. **Nazia Hassan Nisha:** Data curation, Methodology, Software, Visualization. **Arittra Bhattacharjee:** Data curation, Validation, Visualization, Writing – review & editing. **Zeshan Mahmud Chowdhury:** Data curation, Validation, Writing – review & editing. **Ishtiaque Ahammad:** Data curation, Validation, Writing – review & editing. **Keshob Chandra Das:** Data curation, Validation, Writing – review & editing. **Mohammad Shahedur Rahman:** Data curation, Validation, Supervision, Writing – review & editing. **Md. Salimullah:** Conceptualization, Investigation, Resources, Supervision, Writing – review & editing, Data curation, Formal analysis.

