## Supplementary File for "Mapping the PTEN Mutation Landscape: Structural and Functional Drivers of Lung Cancer"

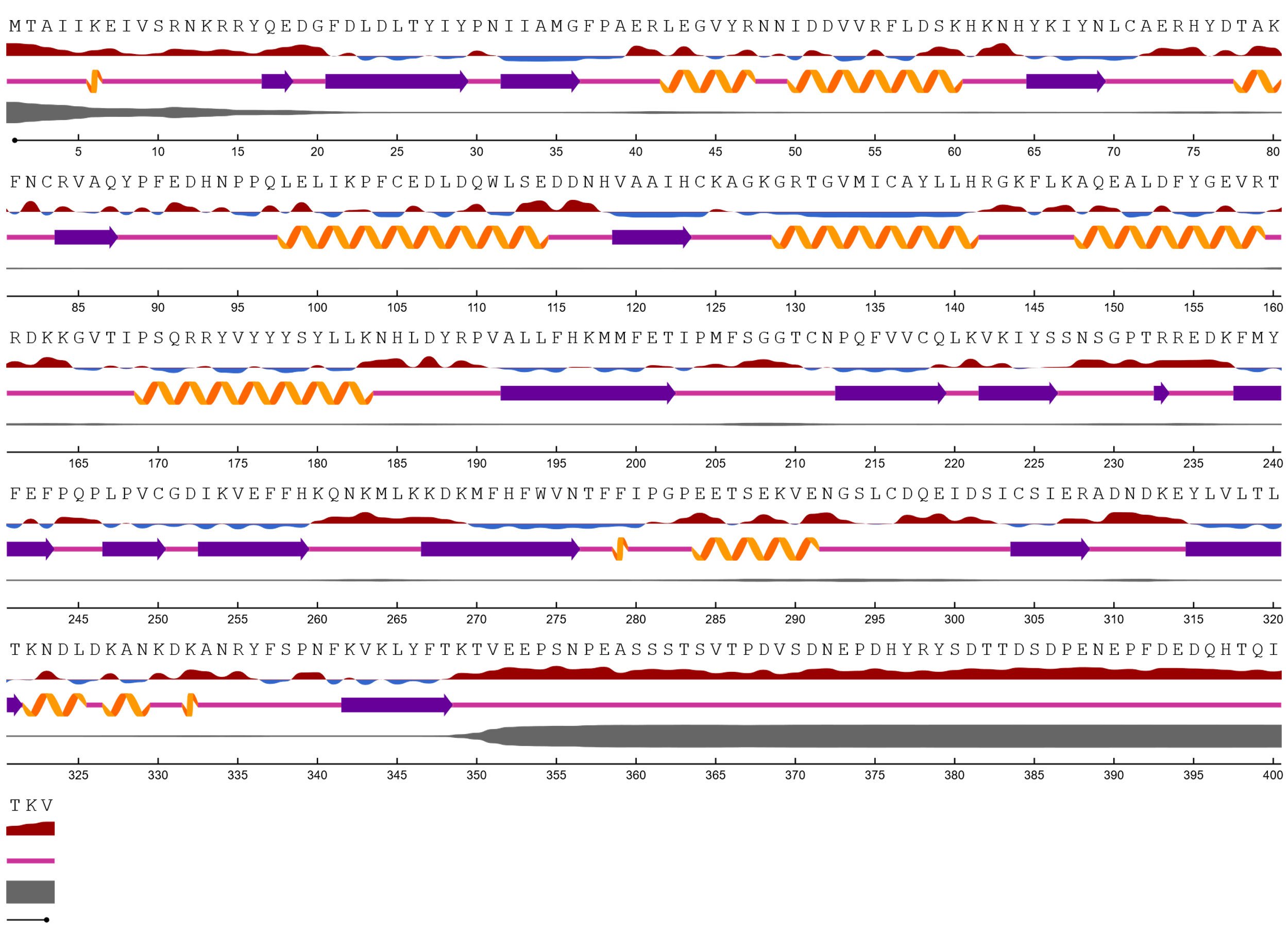

**Figure 1:** NetSurfP-2.0 predicted the PTEN wildtype protein’s surface accessibility, secondary structure and disorders present in different locations.

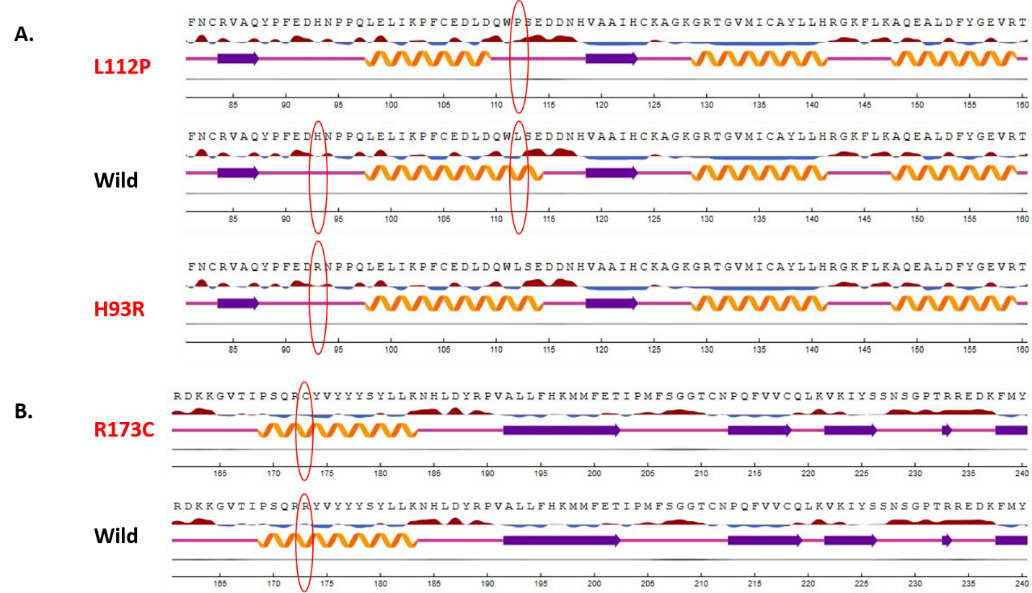

**Figure 2:** NetSurfP-2.0 predicted mutational impact comparison between A) L112P, H93 and wild type, B) R173C and wild type.

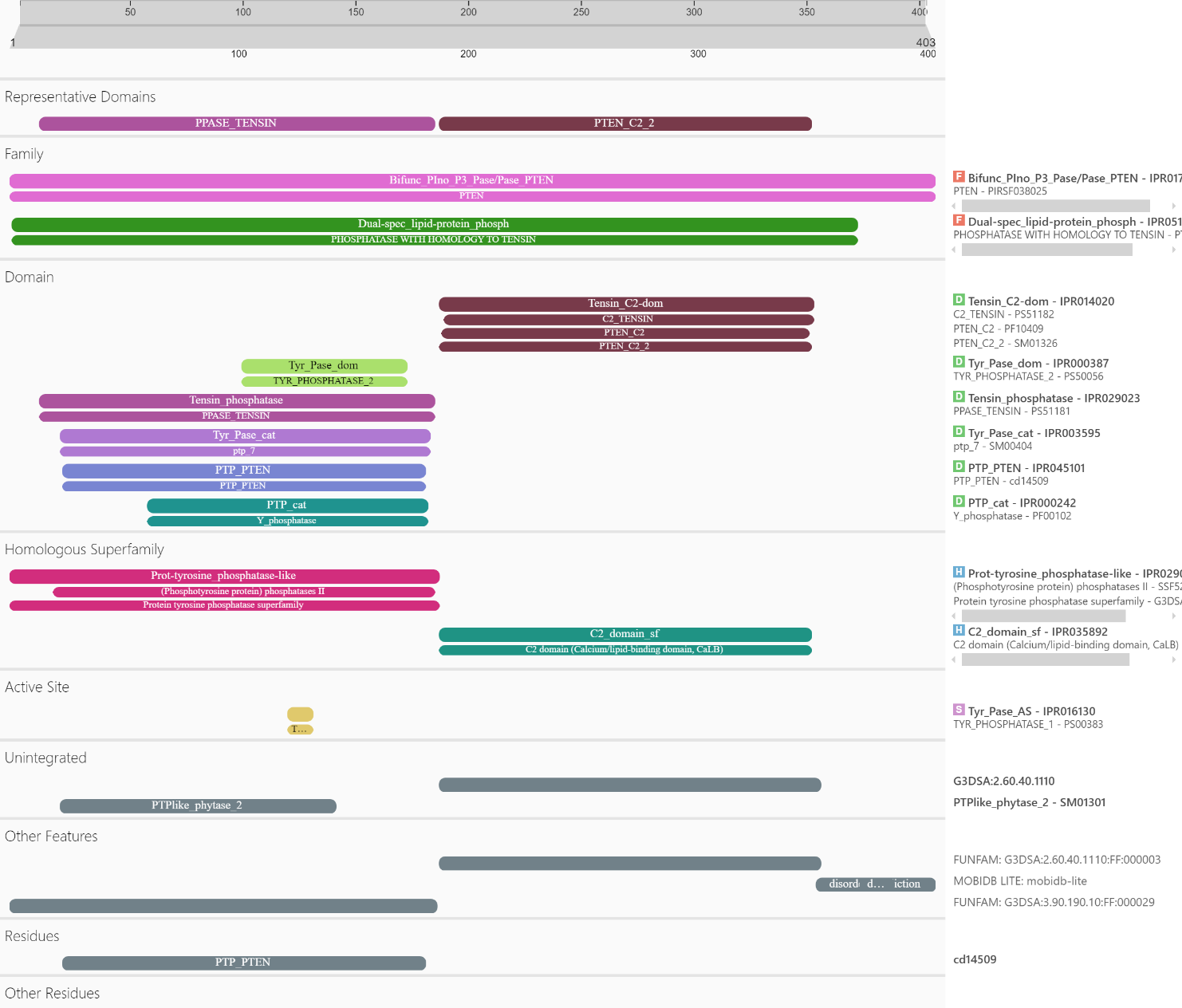

**Figure 3:** Domain, homologous superfamilies, active sites and other features are identified using InterPRO server of PTEN protein.

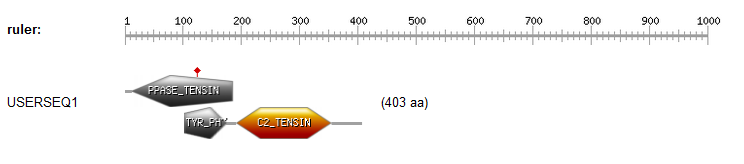

**Figure 4:** PTEN protein domains identified by PROSITE.

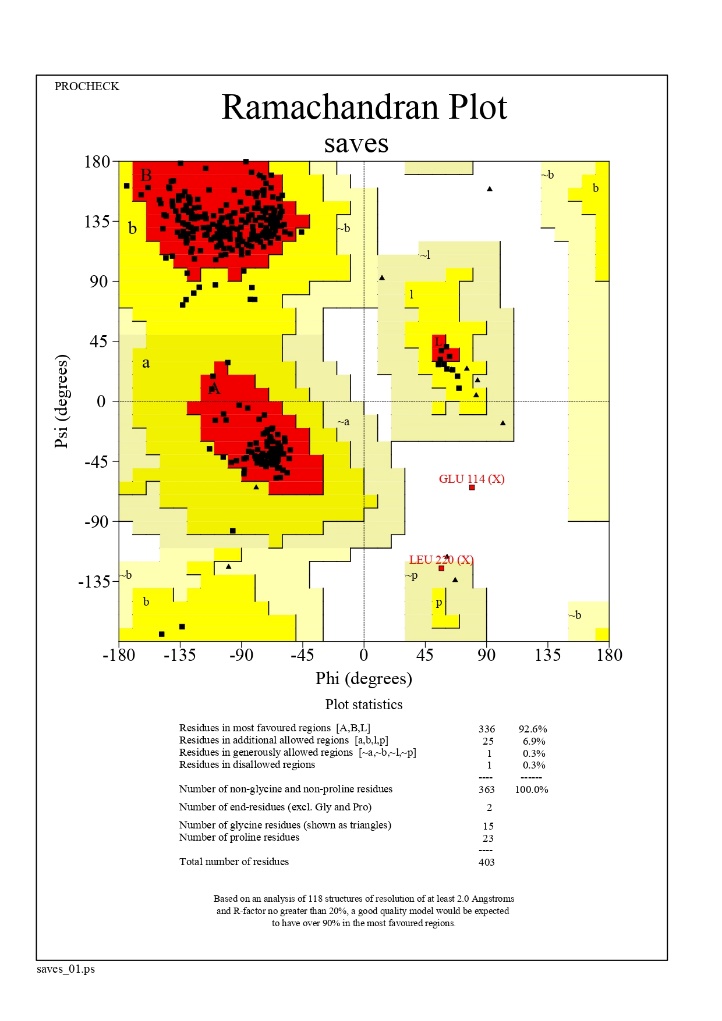

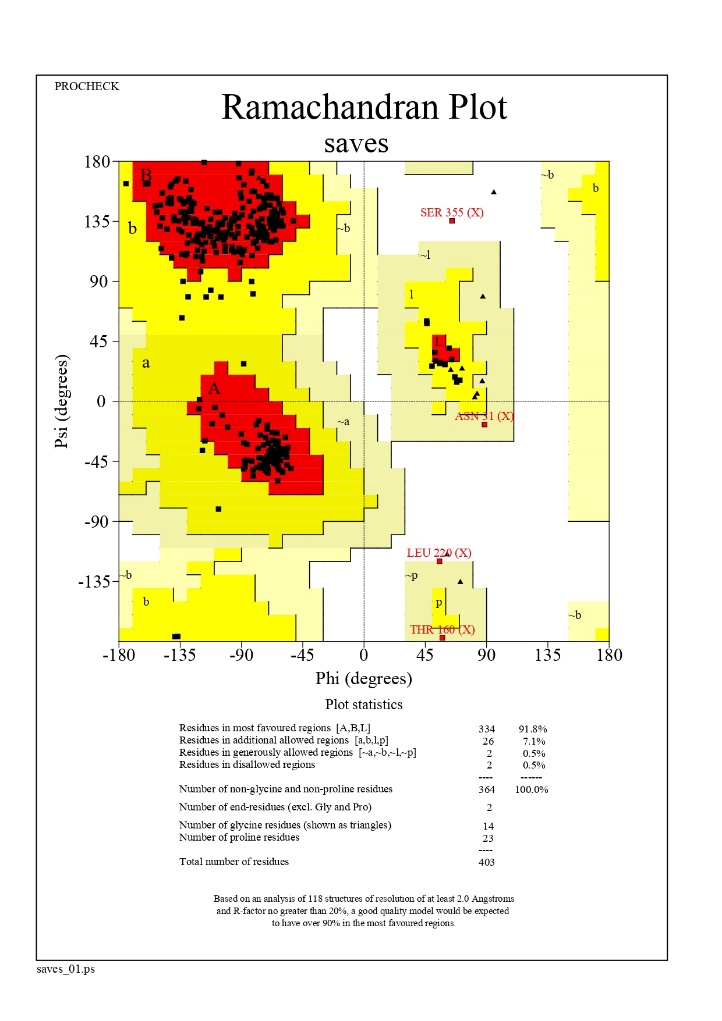

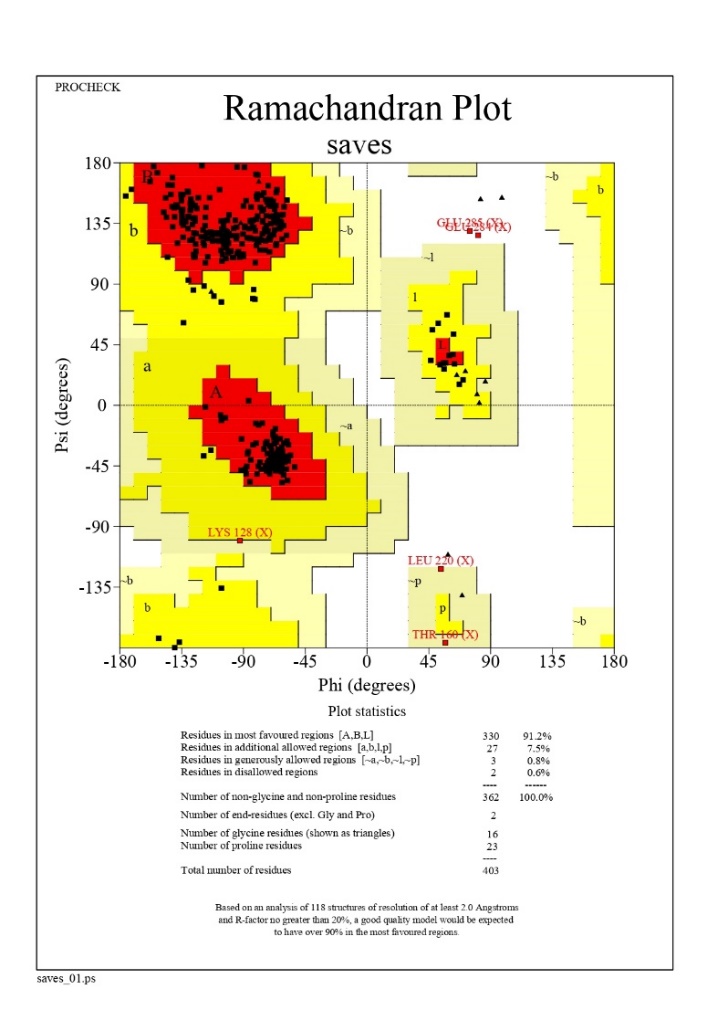

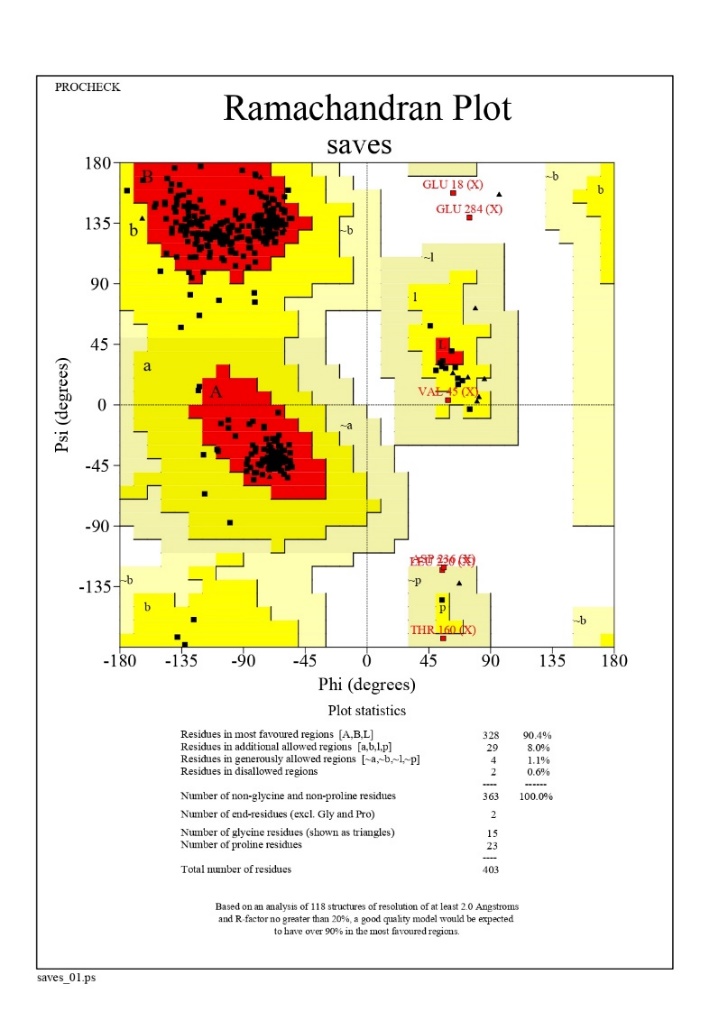

**G132V**

**G129E**

**D252G**

**C124R**

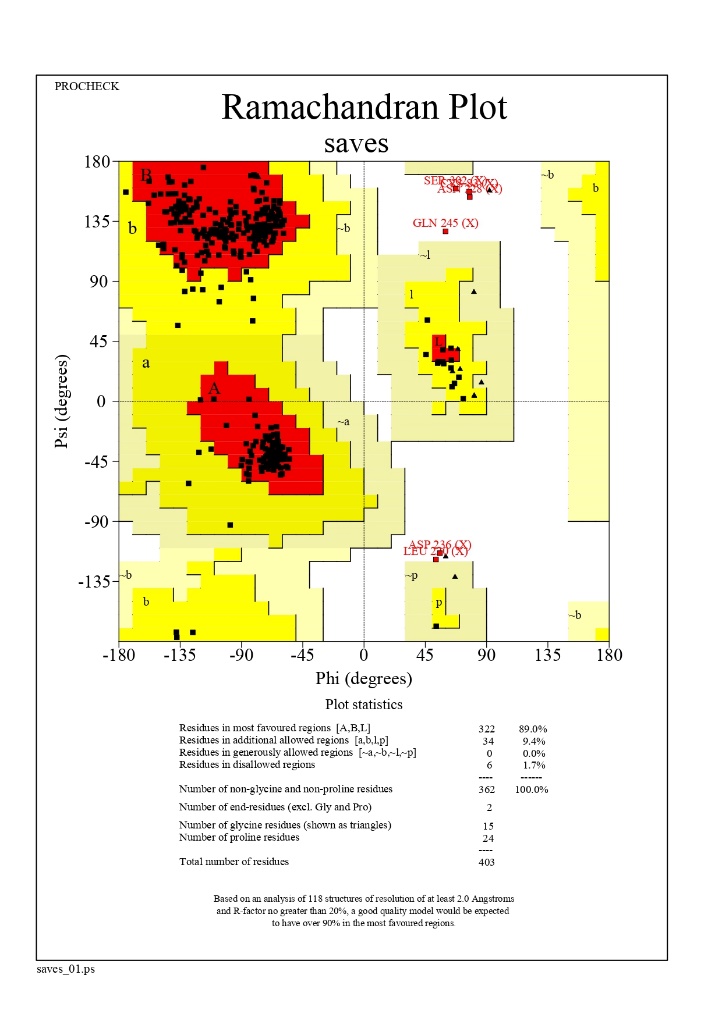

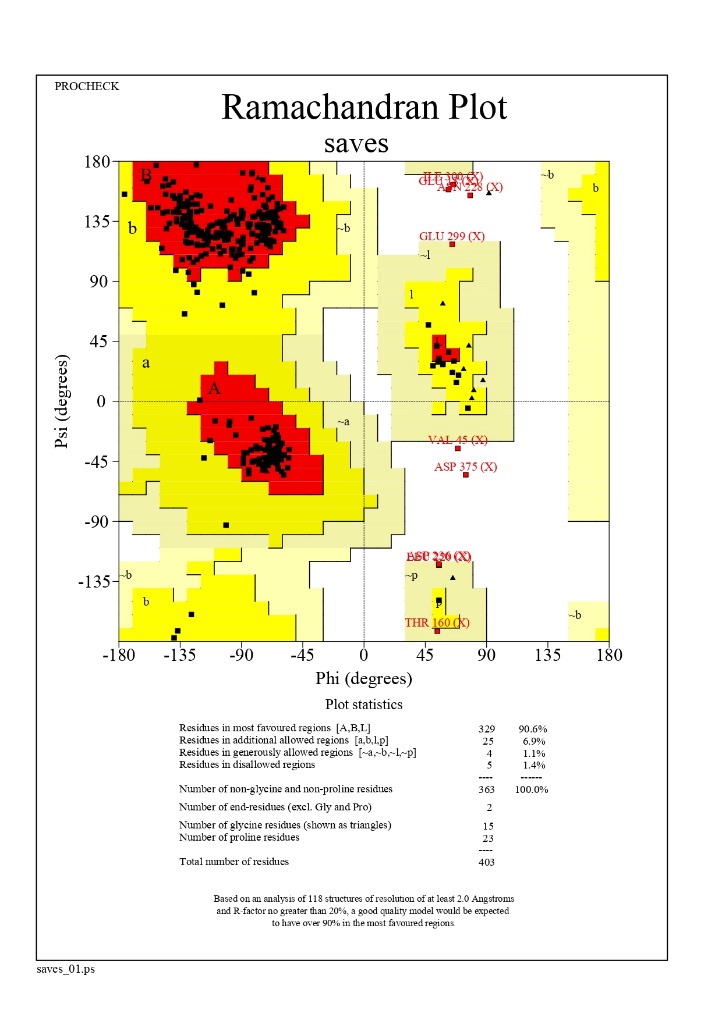

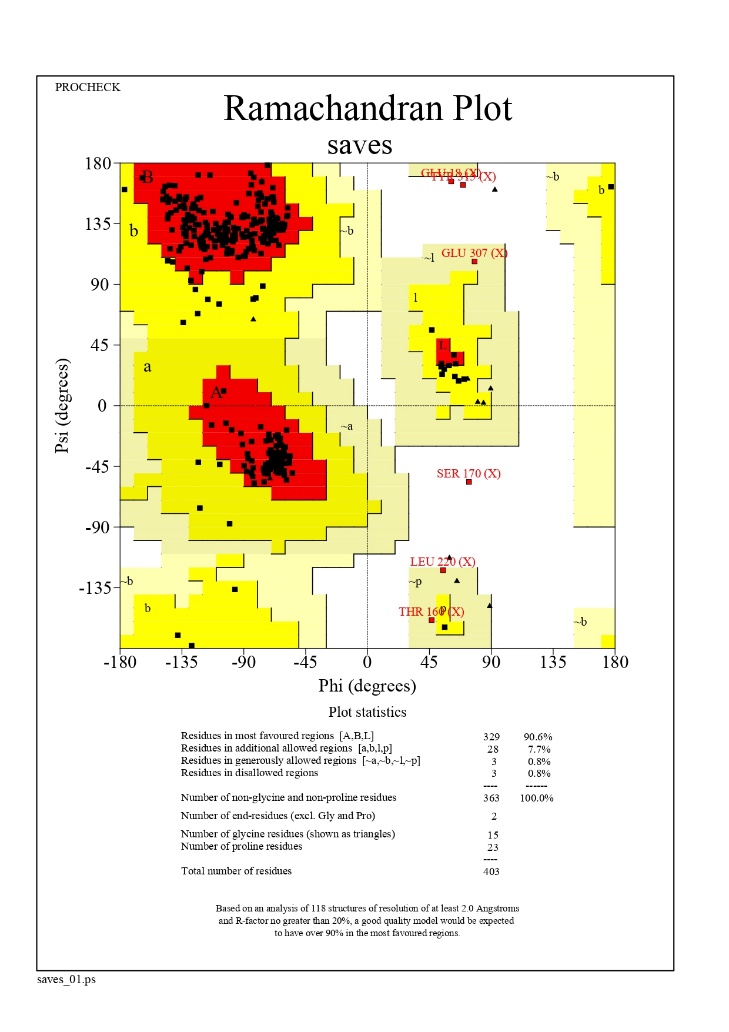

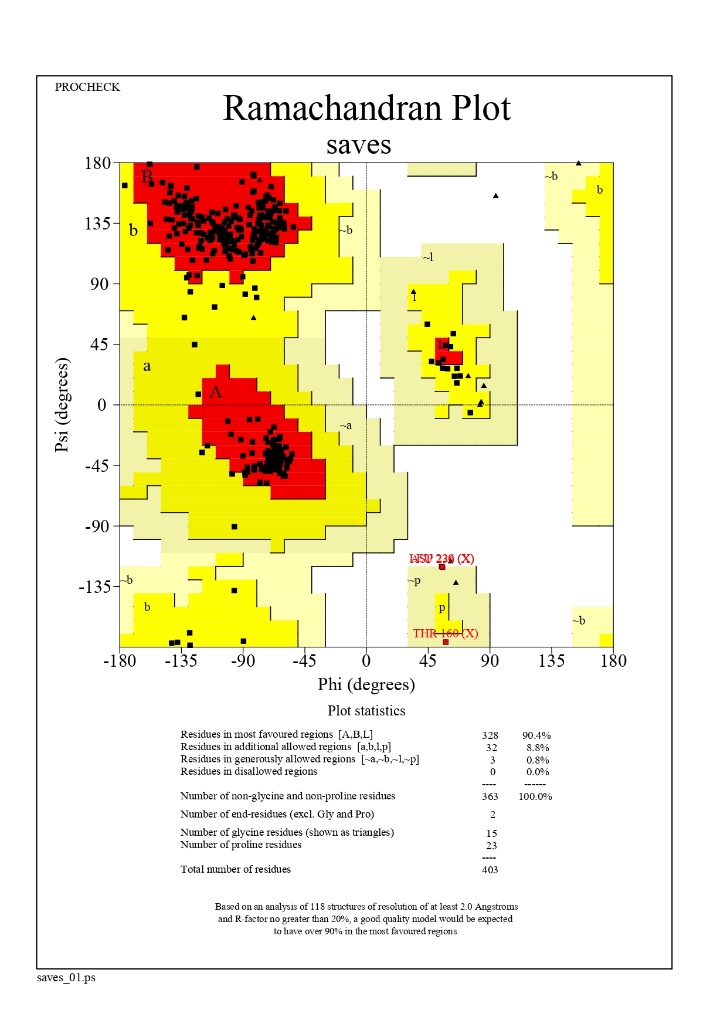

**L70P**

**I135T**

**H123R**

**H93R**

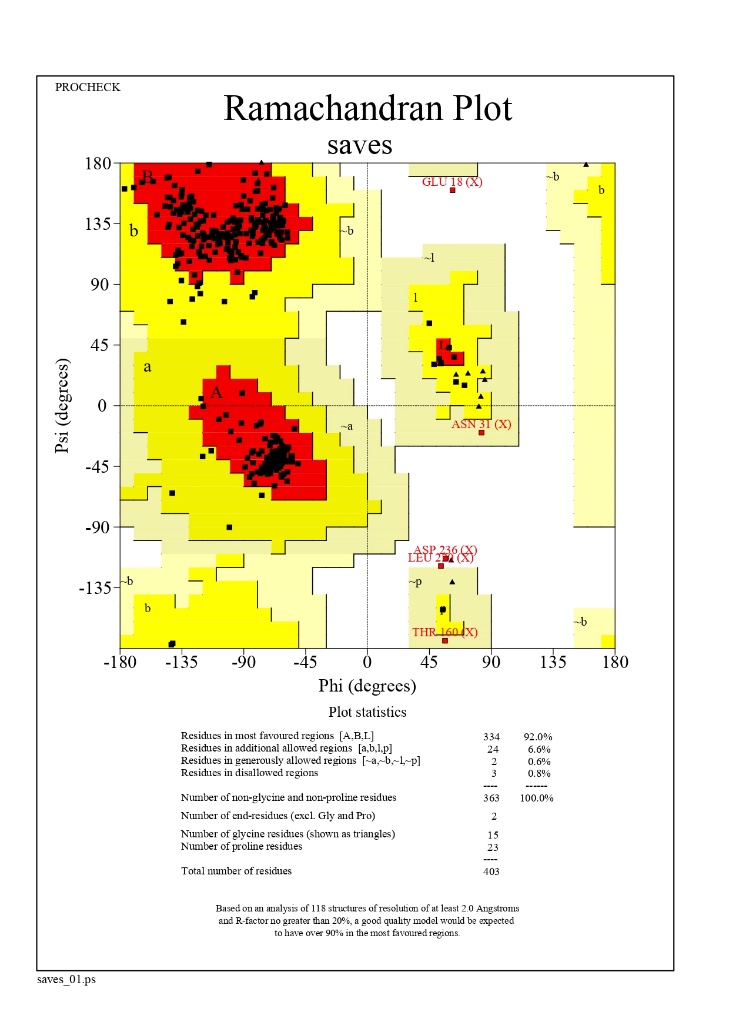

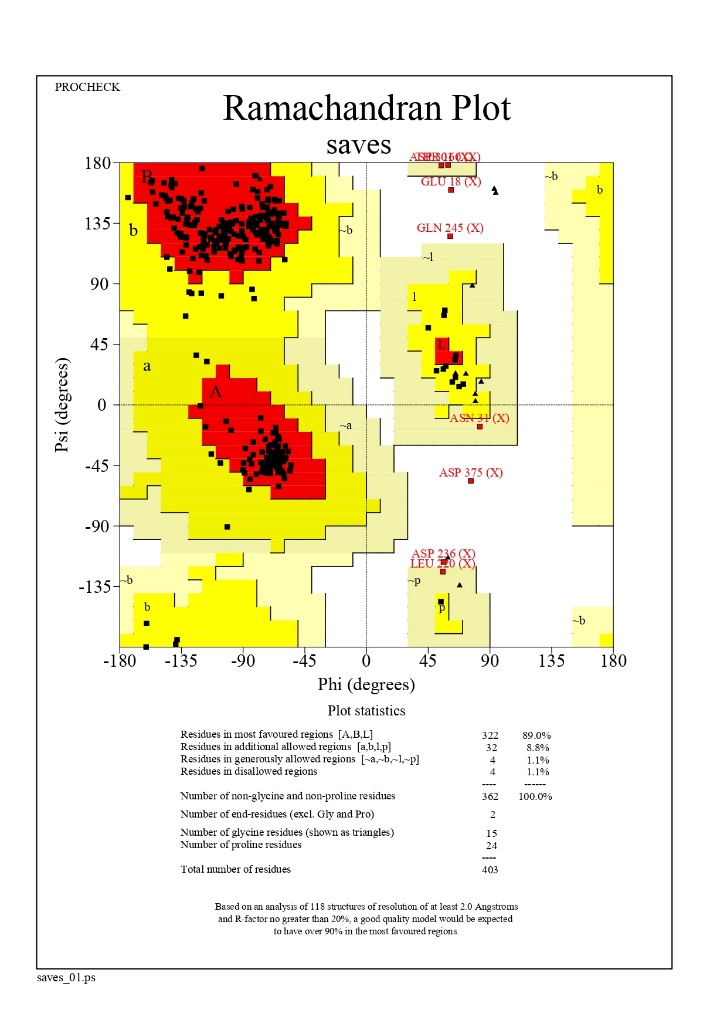

**R173C**

**R130G**

**M35R**

**L112P**

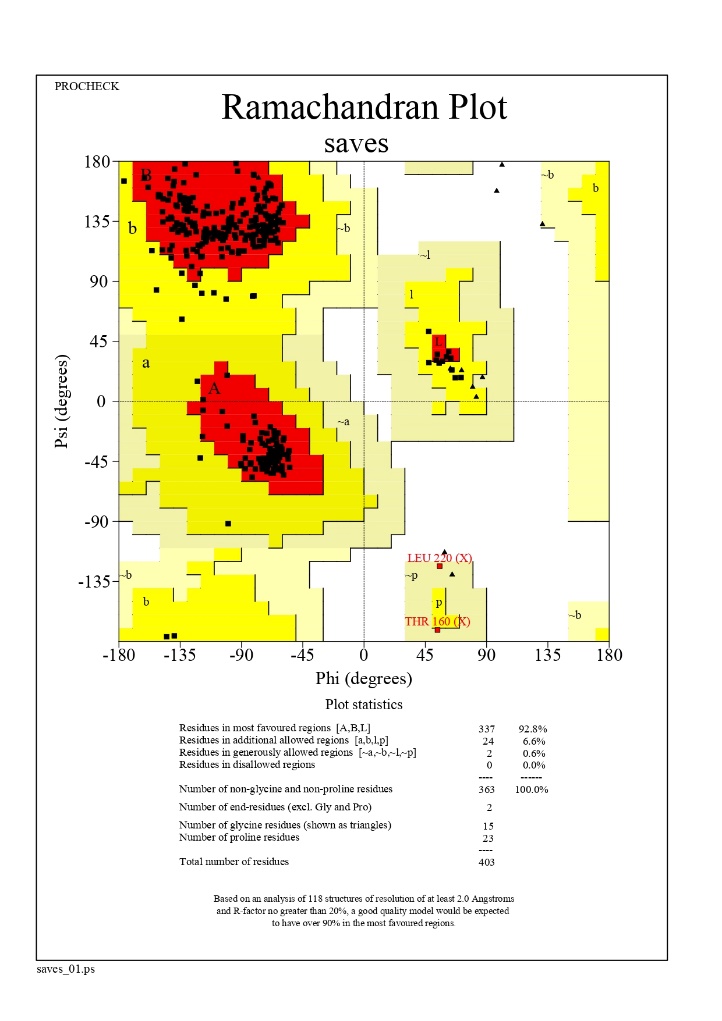

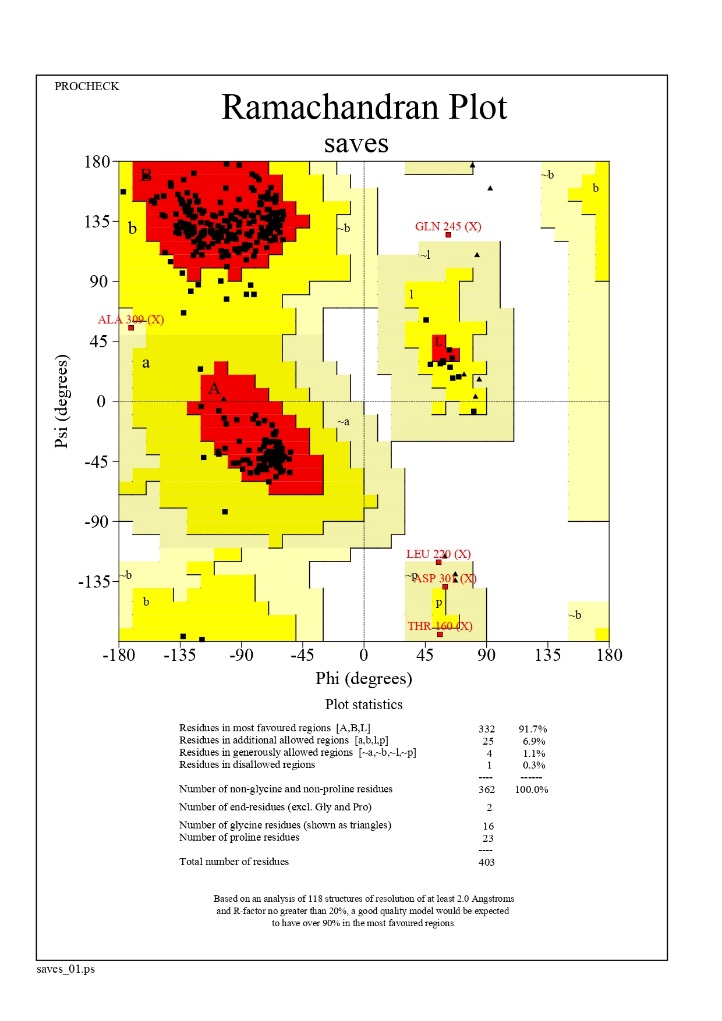

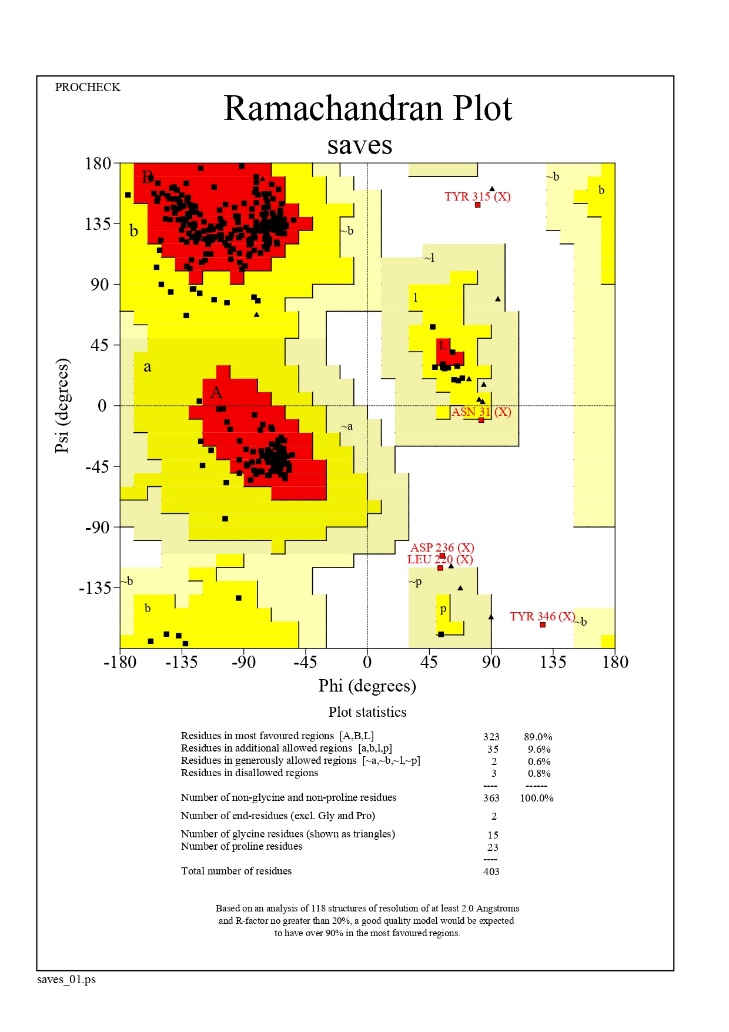

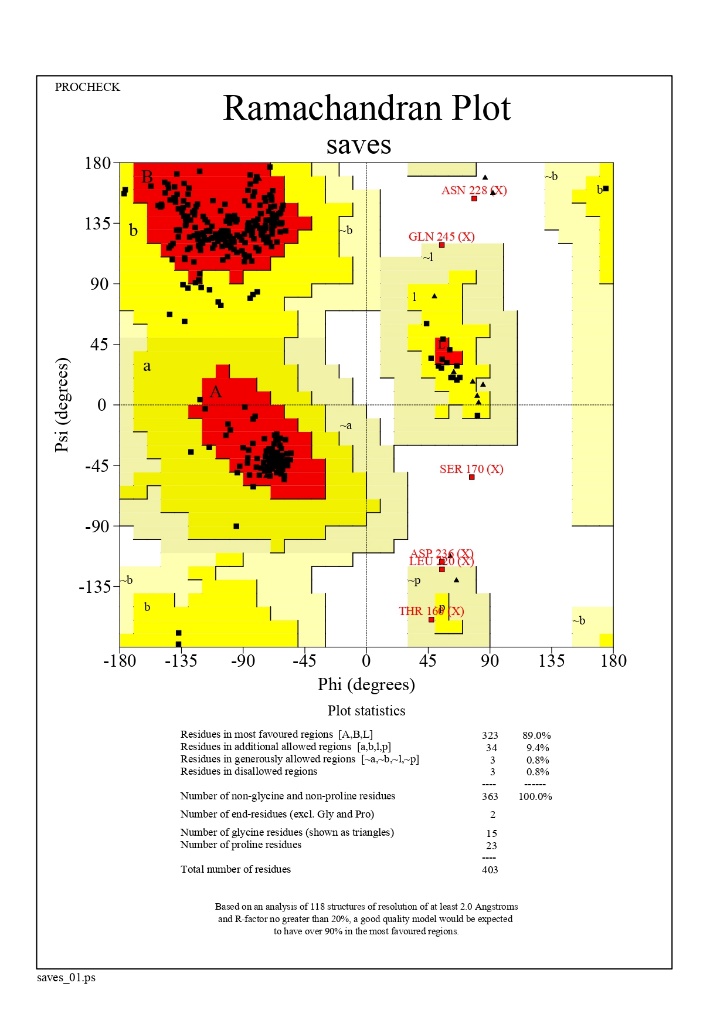

**Wild**

**R130Q**

**V119L**

**R173H**

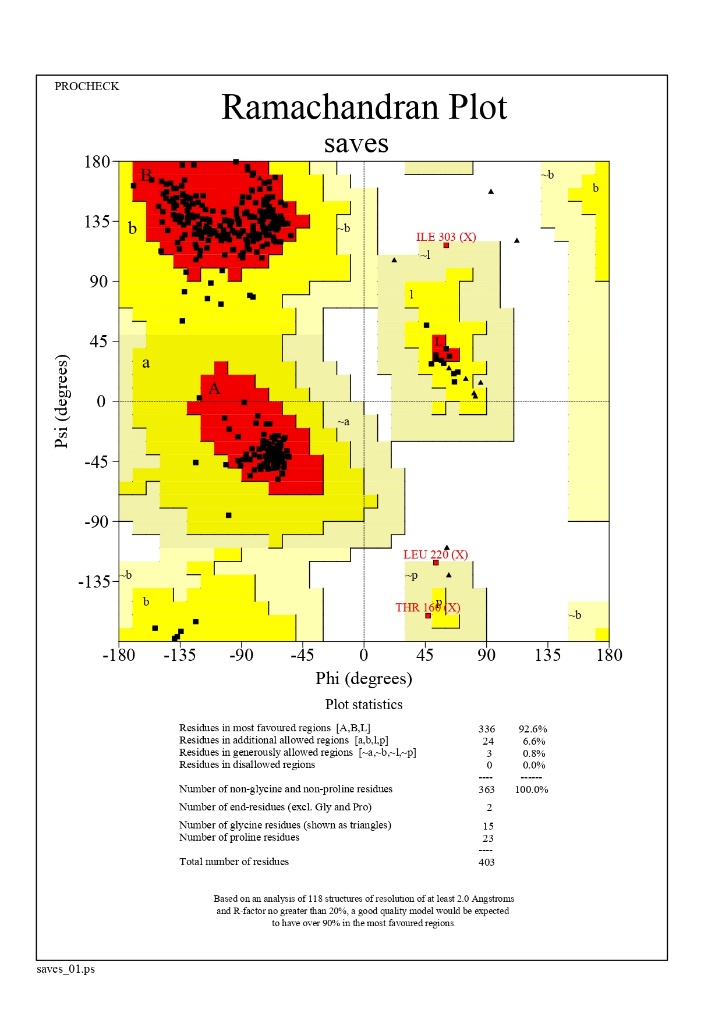

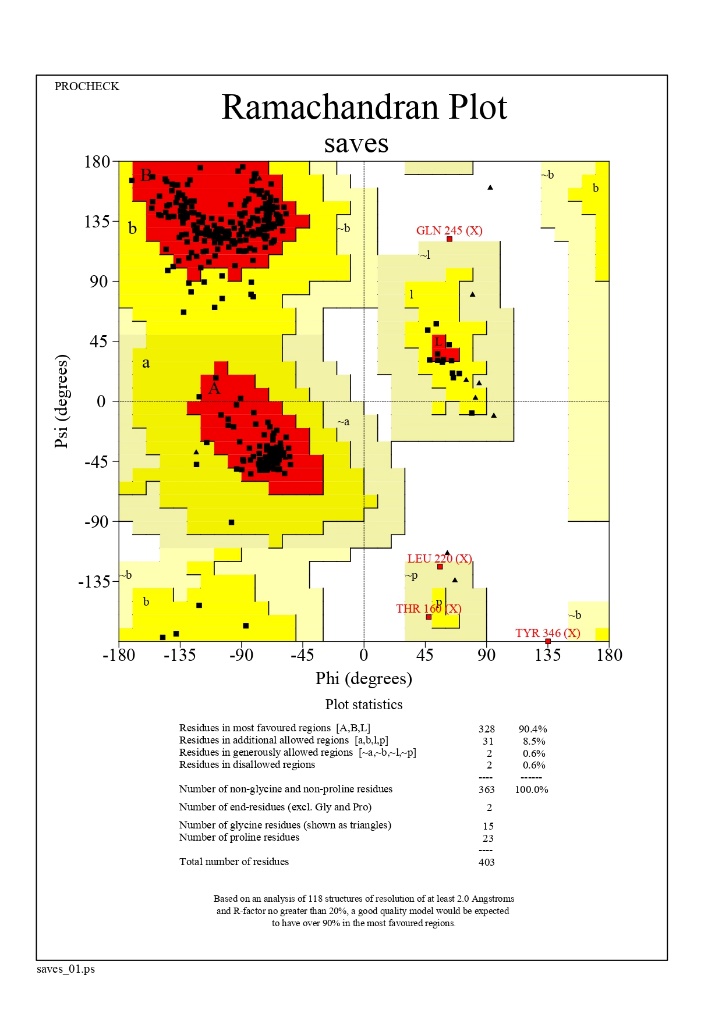

**Figure 5:** Ramachandran plot of PTEN wild type and mutants with the models of GaalxyRefine.

**Table 1:** Structural instability in genes responsible for different types of lung cancer

| Type of lung cancer | Genes involved | References |
| --- | --- | --- |
| Adenocarcinoma of lung | EGFR, ALK, MET, ERBB2, ROS, RET, KRAS, NF1, BRAF, NRAS, PIK3CA, PTEN, AKT1, LKB1, TP53, MDM2, CDKNA2, MYC, SMARCA4, ARID1A, SETD2, NKX2.1/TTF1, KEAP1 | ^1–3^ |
| Squamous cell carcinoma | EGFR, FGFR1-3, ERBB2, ERBB3, DDR2, NF1, KRAS, HRAS, NRAS, RASA1, BRAF, PIK3CA, PTEN, AKT1, AKT2, AKT3, TSC1-2, TP53, CDKNA2, RB1, MLL2, SOX2, TP63, NOTCH1, NOTCH2, ASCL4, FOXP1, KEAP1, NRF2, CUL3 | ^3,4^ |
| Small cell lung carcinoma | FGFR1, PTEN, TP53, RB1, CCNE1, MYC, MYCN, MYCL, EP300, CREBBP, MLL, SLIT2, EPHA7 | ^3,5^ |

**Table 2:** Top 10 in STRING network of adenocarcinoma ranked by degree method

| Rank | Name | Gene Name | Score |
| --- | --- | --- | --- |
| 1 | 9606.ENSP00000256078 | KRAS | 21 |
| 1 | 9606.ENSP00000269305 | TP53 | 21 |
| 1 | 9606.ENSP00000263967 | PIK3CA | 21 |
| 1 | 9606.ENSP00000361021 | PTEN | 21 |
| 5 | 9606.ENSP00000419060 | BRAF | 20 |
| 5 | 9606.ENSP00000478887 | MYC | 20 |
| 5 | 9606.ENSP00000275493 | EGFR | 20 |
| 5 | 9606.ENSP00000351015 | NF1 | 20 |
| 5 | 9606.ENSP00000269571 | ERBB2 | 20 |
| 5 | 9606.ENSP00000358548 | NRAS | 20 |

**Table 3:** Top 10 in STRING network of squamous cell carcinoma ranked by degree method

| Rank | Name | Gene name | Score |
| --- | --- | --- | --- |
| 1 | 9606.ENSP00000269305 | TP53 | 24 |
| 1 | 9606.ENSP00000361021 | PTEN | 24 |
| 3 | 9606.ENSP00000263967 | PIK3CA | 23 |
| 4 | 9606.ENSP00000256078 | KRAS | 22 |
| 4 | 9606.ENSP00000275493 | EGFR | 22 |
| 6 | 9606.ENSP00000498587 | NOTCH1 | 21 |
| 6 | 9606.ENSP00000451828 | AKT1 | 21 |
| 8 | 9606.ENSP00000351015 | NF1 | 20 |
| 9 | 9606.ENSP00000269571 | ERBB2 | 19 |
| 9 | 9606.ENSP00000407586 | HRAS | 19 |

**Table 4:** Top 10 in STRING network of small cell lung carcinoma ranked by degree method

| Rank | Name | Gene Name | Score |
| --- | --- | --- | --- |
| 1 | 9606.ENSP00000361021 | PTEN | 10 |
| 1 | 9606.ENSP00000269305 | TP53 | 10 |
| 3 | 9606.ENSP00000478887 | MYC | 9 |
| 3 | 9606.ENSP00000262367 | CREBBP | 9 |
| 5 | 9606.ENSP00000262643 | CCNE1 | 8 |
| 5 | 9606.ENSP00000281043 | MYCN | 8 |
| 7 | 9606.ENSP00000267163 | RB1 | 7 |
| 7 | 9606.ENSP00000263253 | EP300 | 7 |
| 9 | 9606.ENSP00000436786 | KMT2A | 6 |
| 9 | 9606.ENSP00000393312 | FGFR1 | 6 |

**Table 5:** SIFT prediction of deleterious nsSNPs

| SNP | AA Change | Region | SIFT_Score | SIFT_Prediction |
| --- | --- | --- | --- | --- |
| rs57374291 | D107N | CDS | 0.011 | DELETERIOUS |
| rs121909218 | G129E | CDS | 0 | DELETERIOUS |
| rs121909221 | S170R | CDS | 0.002 | DELETERIOUS |
| rs121909222 | H123R | CDS | 0.002 | DELETERIOUS |
| rs121909223 | C124R | CDS | 0 | DELETERIOUS |
| rs121909224 | R130G | CDS | 0 | DELETERIOUS |
| rs121909225 | M35R | CDS | 0 | DELETERIOUS |
| rs121909226 | L70P | CDS | 0 | DELETERIOUS |
| rs121909229 | R130Q | CDS | 0 | DELETERIOUS |
| rs121909230 | L112P | CDS | 0 | DELETERIOUS |
| rs121909238 | H93R | CDS | 0 | DELETERIOUS |
| rs121909239 | D252G | CDS | 0.01 | DELETERIOUS |
| rs121909241 | G132V | CDS | 0 | DELETERIOUS |
| rs121913293 | R173C | CDS | 0 | DELETERIOUS |
| rs121913294 | R173H | CDS | 0.003 | DELETERIOUS |
| rs139767111 | V119L | CDS | 0.009 | DELETERIOUS |
| rs370795352 | I135T | CDS | 0.002 | DELETERIOUS |

**Table 6:** SNPS&Go prediction of deleterious nsSNPs

| Mutation | Prediction | RI | Probability |
| --- | --- | --- | --- |
| D19N | Disease | 5 | 0.773 |
| M35R | Disease | 9 | 0.941 |
| L70P | Disease | 8 | 0.886 |
| H93R | Disease | 6 | 0.782 |
| L112P | Disease | 8 | 0.881 |
| V119L | Disease | 4 | 0.701 |
| H123R | Disease | 9 | 0.935 |
| C124R | Disease | 9 | 0.964 |
| G129E | Disease | 8 | 0.878 |
| R130G | Disease | 8 | 0.903 |
| R130Q | Disease | 8 | 0.889 |
| G132V | Disease | 8 | 0.922 |
| I135T | Disease | 6 | 0.783 |
| S170R | Disease | 7 | 0.871 |
| R173C | Disease | 8 | 0.92 |
| R173H | Disease | 8 | 0.911 |
| F241S | Disease | 6 | 0.781 |
| D252G | Disease | 8 | 0.905 |

**Table 7:** PMut prediction of deleterious nsSNPs

| Position | Wild type | Mutant | Prediction of disease | Score |
| --- | --- | --- | --- | --- |
| 129 | G | E | TRUE | 0.8034 |
| 170 | S | R | TRUE | 0.9096 |
| 123 | H | R | TRUE | 0.8935 |
| 124 | C | R | TRUE | 0.8941 |
| 130 | R | G | TRUE | 0.8941 |
| 35 | M | R | TRUE | 0.8941 |
| 70 | L | P | TRUE | 0.9034 |
| 130 | R | Q | TRUE | 0.8005 |
| 112 | L | P | TRUE | 0.8941 |
| 93 | H | R | TRUE | 0.7101 |
| 252 | D | G | TRUE | 0.8227 |
| 241 | F | S | TRUE | 0.7791 |
| 132 | G | V | TRUE | 0.9096 |
| 173 | R | C | TRUE | 0.8883 |
| 173 | R | H | TRUE | 0.905 |
| 119 | V | L | TRUE | 0.8005 |
| 135 | I | T | TRUE | 0.7679 |

**Table 8:** PHD_SNP prediction of deleterious nsSNPs

| SNP | State | Score |
| --- | --- | --- |
| D107N | Disease | 7 |
| G129E | Disease | 7 |
| S170R | Disease | 9 |
| H123R | Disease | 8 |
| C124R | Disease | 9 |
| R130G | Disease | 7 |
| M35R | Disease | 9 |
| L70P | Disease | 7 |
| R130Q | Disease | 8 |
| L112P | Disease | 8 |
| D19N | Disease | 1 |
| V217I | Neutral | 9 |
| R234Q | Neutral | 6 |
| H93R | Disease | 6 |
| G132V | Disease | 9 |
| R173C | Disease | 8 |
| R173H | Disease | 7 |
| V119L | Disease | 8 |
| I135T | Disease | 3 |
| Q298E | Neutral | 8 |
| D252G | Disease | 8 |

**Table 9:** Panther prediction of deleterious nsSNPs

| Substitution | Preservation time | Effect | Pdel |
| --- | --- | --- | --- |
| D107N | 1629 | probably damaging | 0.89 |
| G129E | 1629 | probably damaging | 0.89 |
| S170R | 1629 | probably damaging | 0.89 |
| H123R | 1629 | probably damaging | 0.89 |
| C124R | 1629 | probably damaging | 0.89 |
| R130G | 1629 | probably damaging | 0.89 |
| M35R | 1629 | probably damaging | 0.89 |
| L70P | 1629 | probably damaging | 0.89 |
| R130Q | 1629 | probably damaging | 0.89 |
| L112P | 1629 | probably damaging | 0.89 |
| D19N | 1629 | probably damaging | 0.89 |
| V217I | 1629 | probably damaging | 0.89 |
| R234Q | 1629 | probably damaging | 0.89 |
| H93R | 1629 | probably damaging | 0.89 |
| D252G | 1629 | probably damaging | 0.89 |
| F241S | 1629 | probably damaging | 0.89 |
| G132V | 1629 | probably damaging | 0.89 |
| R173C | 1629 | probably damaging | 0.89 |
| R173H | 1629 | probably damaging | 0.89 |
| V119L | 1629 | probably damaging | 0.89 |
| I135T | 1629 | probably damaging | 0.89 |
| Q298E | 1629 | probably damaging | 0.89 |

**Table 10:** Provean prediction of deleterious nsSNPs

| Position | Wild type residue | Mutated residue | Score | Prediction (cutoff=-2.5) | Score | Prediction  (cutoff=0.05) |
| --- | --- | --- | --- | --- | --- | --- |
| 107 | D | N | -4.68 | Deleterious | 0.006 | Damaging |
| 129 | G | E | -7.77 | Deleterious | 0 | Damaging |
| 170 | S | R | -4.58 | Deleterious | 0 | Damaging |
| 123 | H | R | -7.8 | Deleterious | 0 | Damaging |
| 124 | C | R | -11.71 | Deleterious | 0 | Damaging |
| 130 | R | G | -6.75 | Deleterious | 0 | Damaging |
| 35 | M | R | -5.84 | Deleterious | 0.001 | Damaging |
| 70 | L | P | -6.77 | Deleterious | 0 | Damaging |
| 130 | R | Q | -3.86 | Deleterious | 0 | Damaging |
| 112 | L | P | -6.82 | Deleterious | 0 | Damaging |
| 93 | H | R | -7.53 | Deleterious | 0 | Damaging |
| 252 | D | G | -6.49 | Deleterious | 0 | Damaging |
| 132 | G | V | -8.45 | Deleterious | 0 | Damaging |
| 173 | R | C | -7.16 | Deleterious | 0 | Damaging |
| 173 | R | H | -4.14 | Deleterious | 0 | Damaging |
| 119 | V | L | -2.73 | Deleterious | 0.005 | Damaging |
| 135 | I | T | -4.43 | Deleterious | 0.001 | Damaging |

**Table 11:** FATHMM prediction of deleterious nsSNPs

| Substitution | Prediction | Score |
| --- | --- | --- |
| D107N | CANCER | -2.96 |
| G129E | CANCER | -2.99 |
| S170R | CANCER | -6.42 |
| H123R | CANCER | -8.92 |
| C124R | CANCER | -8.81 |
| R130G | CANCER | -5.84 |
| M35R | CANCER | -6.27 |
| L70P | CANCER | -6.5 |
| R130Q | CANCER | -5.84 |
| L112P | CANCER | -6.59 |
| D19N | CANCER | -4.94 |
| V217I | CANCER | -3.38 |
| R234Q | CANCER | -3.33 |
| H93R | CANCER | -6.35 |
| D252G | CANCER | -6.91 |
| F241S | CANCER | -3.41 |
| G132V | CANCER | -7.32 |
| R173C | CANCER | -6.45 |
| R173H | CANCER | -6.42 |
| V119L | CANCER | -6.22 |
| I135T | CANCER | -6.59 |
| Q298E | CANCER | -4.72 |

**Table 12:** I-Mutant 2.0 prediction of protein stability for deleterious nsSNPs of PTEN

| AA Substitution | Effect on Stability | DDG |
| --- | --- | --- |
| G129E | Decrease | -0.2 |
| H123R | Decrease | -1.06 |
| C124R | Decrease | -0.65 |
| R130G | Decrease | -0.45 |
| M35R | Decrease | -1.58 |
| L70P | Decrease | -1.44 |
| R130Q | Decrease | -0.18 |
| L112P | Decrease | -1.82 |
| H93R | Increase | 0.26 |
| D252G | Decrease | -0.28 |
| G132V | Decrease | -0.05 |
| R173C | Decrease | -2.01 |
| R173H | Decrease | -1.8 |
| V119L | Decrease | -0.2 |
| I135T | Decrease | -1.8 |

**Table 13:** Mupro prediction of protein stability for deleterious nsSNPs of PTEN

| SNP | Delta G | Prediction |
| --- | --- | --- |
| G129E | -0.54044838 | DECREASE stability |
| H123R | -0.39016227 | DECREASE stability |
| C124R | -1.212119 | DECREASE stability |
| R130G | -1.3319778 | DECREASE stability |
| M35R | -1.5961575 | DECREASE stability |
| L70P | -2.3639136 | DECREASE stability |
| R130Q | -0.7192958 | DECREASE stability |
| L112P | -1.8174237 | DECREASE stability |
| H93R | -0.3669013 | DECREASE stability |
| D252G | -1.770645 | DECREASE stability |
| G132V | -0.3638809 | DECREASE stability |
| R173C | -1.3272009 | DECREASE stability |
| R173H | -1.854347 | DECREASE stability |
| V119L | -0.28104583 | DECREASE stability |
| I135T | -1.6966794 | DECREASE stability |

**Table 14:** Missense 3D prediction of protein stability for deleterious nsSNPs of PTEN

| **SNP/Properties** | **Disulphide breakage** | **Buried Pro introduced** | **Clash** | **Buried hydrophilic introduced** | **Buried charge introduced** | **Secondary structure altered** | **Buried charge switch** | **Disallowed phi/psi** | **Buried charge replaced** | **Buried Gly replaced** | **Buried H-bond breakage** | **Buried salt bridge breakage** | **Cavity altered** | **Buried / exposed switch** | **Cis pro replaced** | **Gly in a bend** | **Prediction** |
| --- | --- | --- | --- | --- | --- | --- | --- | --- | --- | --- | --- | --- | --- | --- | --- | --- | --- |
| **G129E** | N | N | N | N | N | N | N | N | N | N | N | N | N | N | N | N | **Neutral** |
| **H123R** | N | N | N | N | N | N | N | N | N | N | **Y** | N | N | N | N | N | **Damaging** |
| **C124R** | N | N | N | **Y** | **Y** | N | N | N | N | N | N | N | N | N | N | N | **Damaging** |
| **R130G** | N | N | N | N | N | N | N | N | N | N | N | **W** | **Y** | N | N | N | **Damaging** |
| **M35R** | N | N | N | **Y** | **Y** | N | N | N | N | N | N | N | N | N | N | N | **Damaging** |
| **L70P** | N | **Y** | N | N | N | N | N | N | N | N | N | N | N | N | N | N | **Damaging** |
| **R130Q** | N | N | N | N | N | N | N | N | **Y** | N | N | **W** | N | N | N | N | **Damaging** |
| **L112P** | N | N | N | N | N | N | N | N | N | N | N | N | N | N | N | N | **Neutral** |
| **H93R** | N | N | N | N | N | N | N | N | N | N | N | N | **Y** | N | N | N | **Damaging** |
| **D252G** | N | N | N | N | N | N | N | N | N | N | **Y** | N | N | **Y** | N | N | **Damaging** |
| **G132V** | N | N | N | N | N | N | N | N | N | **Y** | N | N | **E** | N | N | N | **Damaging** |
| **R173C** | N | N | N | N | N | N | N | N | N | N | N | N | N | N | N | N | **Neutral** |
| **R173H** | N | N | N | N | N | N | N | N | N | N | N | N | N | N | N | N | **Neutral** |
| **V119L** | N | N | N | N | N | N | N | N | N | N | N | N | **E** | N | N | N | **Neutral** |
| **I135T** | N | N | N | N | N | N | N | N | N | N | N | N | **E** | N | N | N | **Neutral** |

**Table 15:** Hope server predicted protein structural stability of native and mutant PTEN proteins

| **SNP** | **Wild type** | **Mutant type** | **Properties** |
| --- | --- | --- | --- |
| **G129E** | 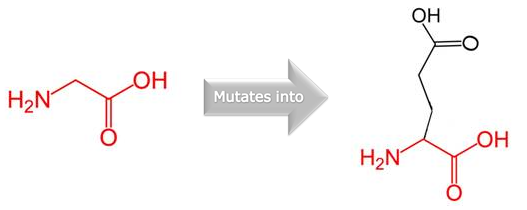 | 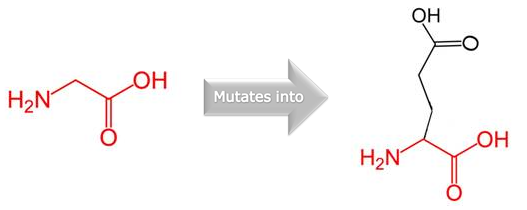 | - The mutant residue is bigger than the wild-type residue, this might lead to bumps. - The wild-type residue charge was NEUTRAL, the mutant residue charge is NEGATIVE. - The wild-type residue is more hydrophobic than the mutant residue. - The mutation introduces a charge, this can cause repulsion of ligands or other residues with the same charge. - The torsion angles for this residue are unusual. only glycine is flexible enough to make these torsion angles, mutation into another residue will force the local backbone into an incorrect conformation and will disturb the local structure. - The mutation is located within a domain, annotated in UniProt as: Phosphatase tensin-type |
| **H123R** | 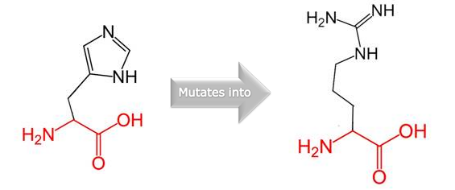 | 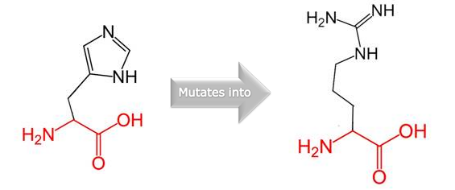 | - The mutant residue is bigger than the wild-type residue. - The wild-type residue charge was NEUTRAL, the mutant residue charge is POSITIVE. - The mutation is located within a domain, annotated in UniProt as: Phosphatase tensin-type - There is a difference in charge between the wild-type and mutant amino acid. - The mutation introduces a charge, this can cause repulsion of ligands or other residues with the same charge. - The wild-type and mutant amino acids differ in size. - The mutant residue is bigger, this might lead to bumps. |
| **C124R** | 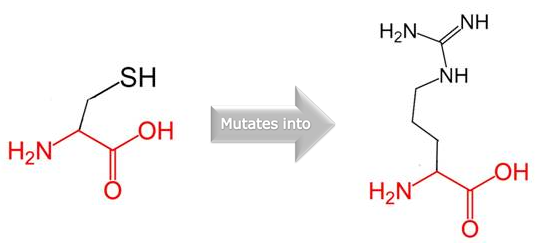 | 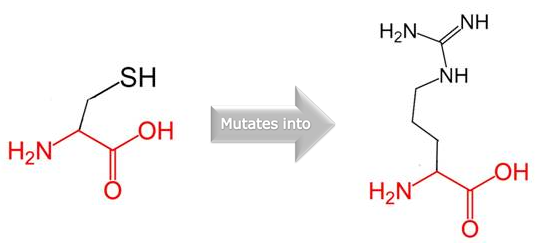 | - The mutant residue is bigger than the wild-type residue. - The wild-type residue charge was NEUTRAL, the mutant residue charge is POSITIVE. - The wild-type residue is more hydrophobic than the mutant residue. - The mutation is located within a domain, annotated in UniProt as: Phosphatase tensin-type - There is a difference in charge between the wild-type and mutant amino acid. - The mutation introduces a charge, this can cause repulsion of ligands or other residues with the same charge. - The wild-type and mutant amino acids differ in size. - The mutant residue is bigger, this might lead to bumps. - The hydrophobicity of the wild-type and mutant residue differs. Hydrophobic interactions, either in the core of the protein or on the surface, will be lost. |
| **R130G** | 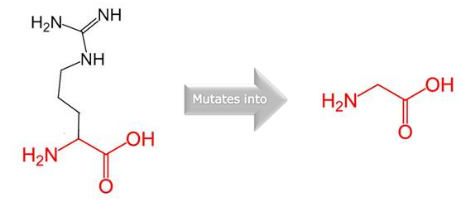 | 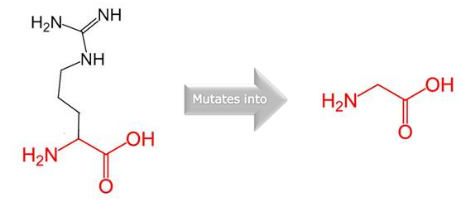 | - The mutant residue is smaller than the wild-type residue. - The wild-type residue charge was POSITIVE, the mutant residue charge is NEUTRAL. - The mutant residue is more hydrophobic than the wild-type residue. - The mutation is located within a domain, annotated in UniProt as: Phosphatase tensin-type - There is a difference in charge between the wild-type and mutant amino acid. - The charge of the wild-type residue will be lost, this can cause loss of interactions with other molecules or residues. - The mutant residue is smaller, this might lead to loss of interactions. - The hydrophobicity of the wild-type and mutant residue differs. The mutation introduces a more hydrophobic residue at this position. This can result in loss of hydrogen bonds and/or disturb correct folding. |
| **M35R** | 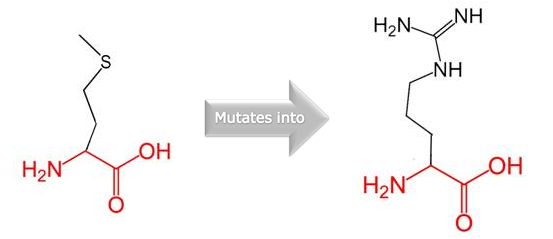 | 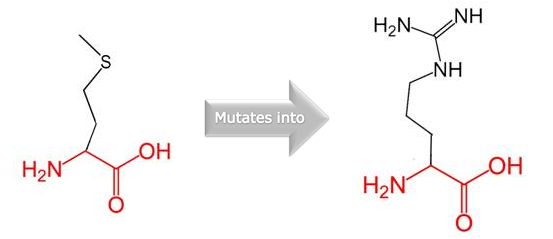 | - The mutant residue is bigger than the wild-type residue. - The wild-type residue charge was NEUTRAL, the mutant residue charge is POSITIVE. - The wild-type residue is more hydrophobic than the mutant residue. - The mutation is located within a domain, annotated in UniProt as: Phosphatase tensin-type - There is a difference in charge between the wild-type and mutant amino acid. - The mutation introduces a charge, this can cause repulsion of ligands or other residues with the same charge. - The wild-type and mutant amino acids differ in size. - The mutant residue is bigger, this might lead to bumps. - The hydrophobicity of the wild-type and mutant residue differs. Hydrophobic interactions, either in the core of the protein or on the surface, will be lost. |
| **L70P** | 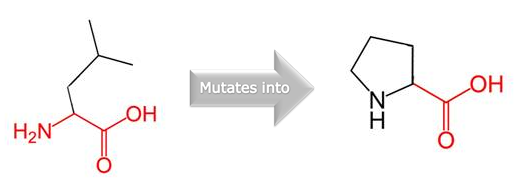 | 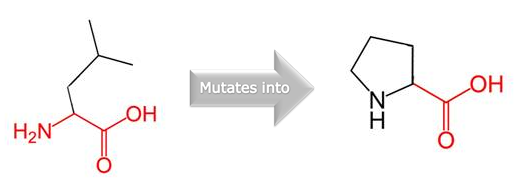 | - The mutation is located within a domain, annotated in UniProt as: Phosphatase tensin-type - The wild-type and mutant amino acids differ in size. - The mutant residue is smaller, this might lead to loss of interactions. - The wild-type and mutant amino acids differ in size. - The mutant residue is smaller than the wild-type residue. - The mutation will cause an empty space in the core of the protein. |
| **R130Q** | 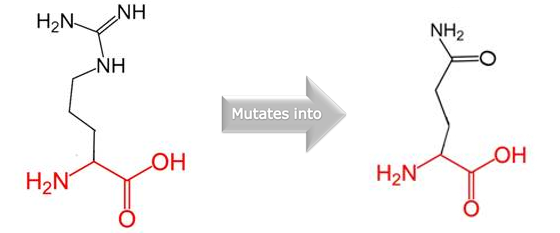 | 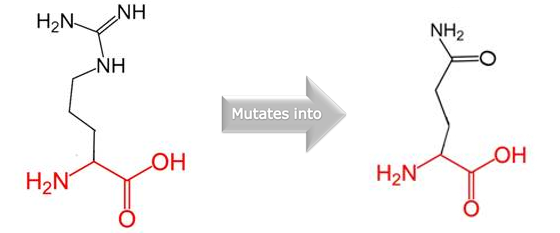 | - The mutant residue is smaller than the wild-type residue. - The wild-type residue charge was POSITIVE, the mutant residue charge is NEUTRAL. - The mutation is located within a domain, annotated in UniProt as: Phosphatase tensin-type - There is a difference in charge between the wild-type and mutant amino acid. - The charge of the wild-type residue will be lost, this can cause loss of interactions with other molecules or residues. |
| **L112P** | 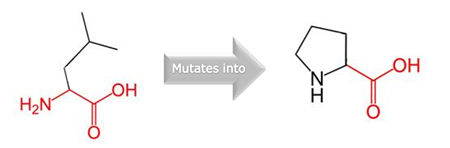 | 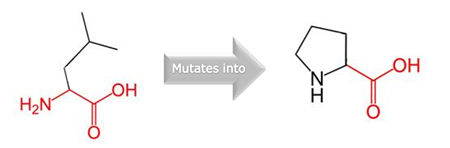 | - The mutant residue is smaller than the wild-type residue. - The mutation is located within a domain, annotated in UniProt as: Phosphatase tensin-type - The mutation introduces an amino acid with different properties, which can disturb this domain and abolish its function. - The wild-type residue is predicted (using the Reprof software) to be located in an α-helix. - Proline disrupts an α-helix when not located at one of the first 3 positions of that helix. In case of the mutation at hand, the helix will be disturbed and this can have severe effects on the structure of the protein. |
| **H93R** | 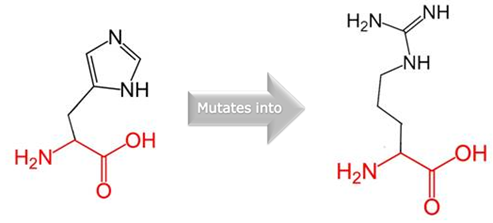 | 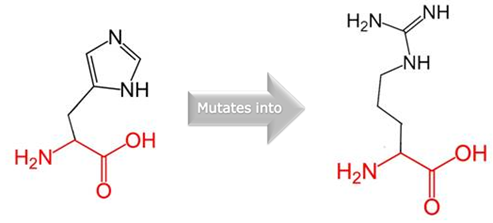 | - The mutant residue is bigger than the wild-type residue. - The wild-type residue charge was NEUTRAL, the mutant residue charge is POSITIVE. - The mutation is located within a domain, annotated in UniProt as: Phosphatase tensin-type - The mutation introduces a charge, this can cause repulsion of ligands or other residues with the same charge. - The wild-type and mutant amino acids differ in size. The mutant residue is bigger, this might lead to bumps. |
| **D252G** | 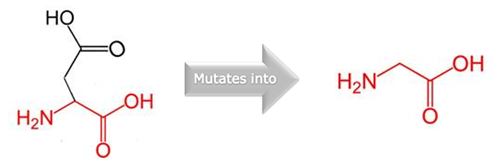 | 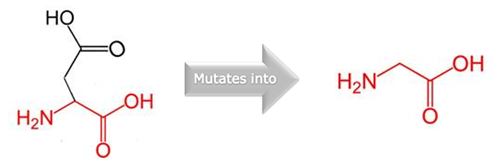 | - The mutant residue is smaller than the wild-type residue. - The wild-type residue charge was NEGATIVE, the mutant residue charge is NEUTRAL. - The mutant residue is more hydrophobic than the wild-type residue. - The mutation is located within a domain, annotated in UniProt as: C2 tensin-type - The charge of the wild-type residue will be lost, this can cause loss of interactions with other molecules or residues.The mutant residue is smaller, this might lead to loss of interactions. - The hydrophobicity of the wild-type and mutant residue differs. The mutation introduces a more hydrophobic residue at this position. This can result in loss of hydrogen bonds and/or disturb correct folding. |
| **G132V** | 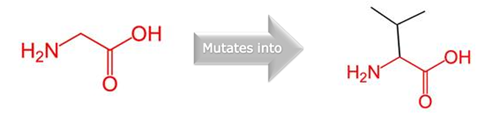 | 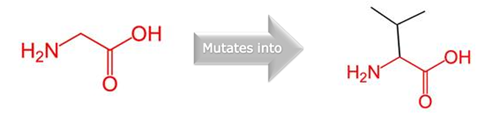 | - The mutant residue is bigger than the wild-type residue. - The mutant residue is more hydrophobic than the wild-type residue. - The mutation is located within a domain, annotated in UniProt as: Phosphatase tensin-type - The wild-type and mutant amino acids differ in size. - The mutant residue is bigger, this might lead to bumps. - The torsion angles for this residue are unusual. only glycine is flexible enough to make these torsion angles, mutation into another residue will force the local backbone into an incorrect conformation and will disturb the local structure. |
| **R173C** | 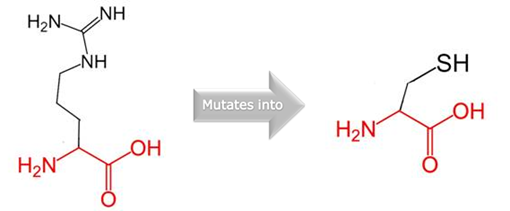 | 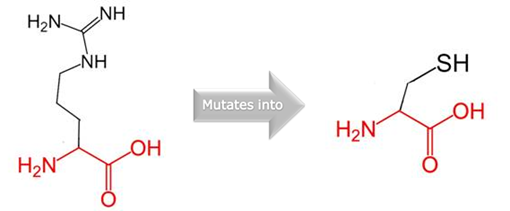 | - The mutant residue is smaller than the wild-type residue. - The wild-type residue charge was POSITIVE, the mutant residue charge is NEUTRAL. - The mutant residue is more hydrophobic than the wild-type residue. - The mutation is located within a domain, annotated in UniProt as: Phosphatase tensin-type - The charge of the wild-type residue will be lost, this can cause loss of interactions with other molecules or residues. - The wild-type and mutant amino acids differ in size. - The mutant residue is smaller, this might lead to loss of interactions. - The hydrophobicity of the wild-type and mutant residue differs. The mutation introduces a more hydrophobic residue at this position. This can result in loss of hydrogen bonds and/or disturb correct folding. |
| **R173H** | 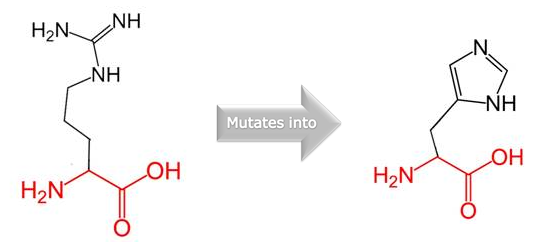 | 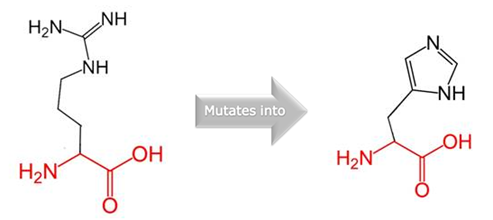 | - The mutant residue is smaller than the wild-type residue. - The wild-type residue charge was POSITIVE, the mutant residue charge is NEUTRAL. - The mutation is located within a domain, annotated in UniProt as: Phosphatase tensin-type - The charge of the wild-type residue will be lost, this can cause loss of interactions with other molecules or residues. - The smaller size of mutant residue may lead to loss of interactions. |
| **V119L** | 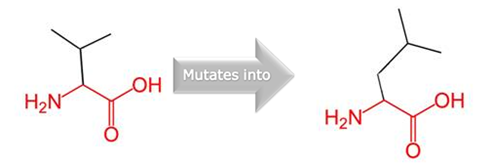 | 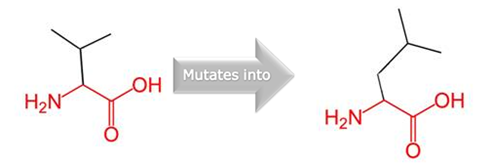 | - The mutant residue is bigger than the wild-type residue. - The mutation is located within a domain, annotated in UniProt as: Phosphatase tensin-type - The wild-type and mutant amino acids differ in size.The mutant residue is bigger, this might lead to bumps. |
| **I135T** | 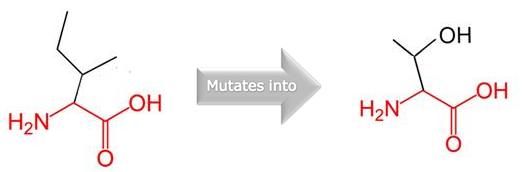 | 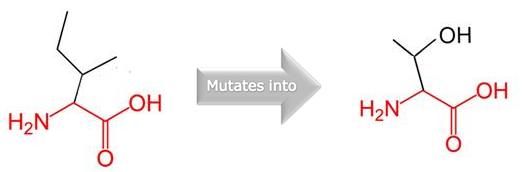 | - The mutant residue is smaller than the wild-type residue. - The wild-type residue is more hydrophobic than the mutant residue. - The mutation is located within a domain, annotated in UniProt as: Phosphatase tensin-type - The wild-type and mutant amino acids differ in size. - The mutant residue is smaller, this might lead to loss of interactions. - The hydrophobicity of the wild-type and mutant residue differs. Hydrophobic interactions, either in the core of the protein or on the surface, will be lost. |

**Table 16:** CASTp predicted biggest pockets for domain regions of PTEN variants

| **SNP** | **Pockets selected** | **Area (SA) Å2** | **Volume (SA) Å3** |
| --- | --- | --- | --- |
| C124R | 1 | 1979.307 | 3523.926 |
| D252G | 1 | 1293.534 | 756.275 |
| G129E | 1,2 | 865.102, 504.847 | 961.797, 817.721 |
| G132V | 1 | 1452.634 | 890.018 |
| H93R | 1,3 | 686.856, 340.168 | 1591.007, 220.743 |
| H123R | 1 | 2522.38 | 2629.123 |
| I135T | 1 | 1876.192 | 2271.902 |
| L70P | 1 | 1722.223 | 1866.674 |
| L112P | 1 | 2316.626 | 2948.699 |
| M35R | 1 | 2684.376 | 5712.732 |
| R130G | 1,2 | 564.033, 1023.374 | 2373.778, 1231.474 |
| R130Q | 1,2 | 205.220, 624.757 | 863.008, 575.151 |
| R173C | 1 | 1614.59 | 1754.777 |
| R173H | 1 | 2120.846 | 2301.925 |
| V119L | 1 | 1101.047 | 2253.884 |
| Wild | 1,2 | 379.486, 535.836 | 2192.634, 1191.744 |

**Table 17:** DynaMut server predicted stability nature of mutant PTEN proteins

| Mutation | Stability (ΔΔG) | Impact |
| --- | --- | --- |
| C124R | -0.271 kcal/mol | Destabilizing |
| D252G | - 0.041 kcal/mol | Destabilizing |
| G129E | 0.097 kcal/mol | Stabilizing |
| G132V | 0.117 kcal/mol | Stabilizing |
| H93R | 0.875 kcal/mol | Stabilizing |
| H123R | - 0.663 kcal/mol | Destabilizing |
| I135T | - 1.708 kcal/mol | Destabilizing |
| L70P | - 2.116 kcal/mol | Destabilizing |
| L112P | - 1.858 kcal/mol | Destabilizing |
| M35R | - 0.882 kcal/mol | Destabilizing |
| R130G | - 0.597 kcal/mol | Destabilizing |
| R130Q | 0.499 kcal/mol | Stabilizing |
| R173C | - 2.493 kcal/mol | Destabilizing |
| R173H | - 1.173 kcal/mol | Destabilizing |
| V119L | 0.420 kcal/mol | Stabilizing |

**Table 18:** Presence of deleterious SNPs on InterPro predicted domains, their ranges and activities

| Domain Name | Domain Range | Activity | Number of deleterious SNPs present | Entry | Reference |
| --- | --- | --- | --- | --- | --- |
| Tensin phosphatase, C2 domain | 188-350 | Can interact with phospholipid membranes in a Ca^2+^ independent manner | 1 | **InterPro:** IPR014020 | ^6^ |
|  |  |  |  | **SMART:**  SM01326 |  |
|  |  |  |  | **PROSITE profiles:**  PS51182 |  |
|  |  |  |  | **Pfam:**  PF10409 |  |
| Tyrosine-specific protein phosphatases domain | 102-173 | Removal of phosphate group linked with tyrosine | 11 | **InterPro:** IPR014020 | ^7^ |
|  |  |  |  | **PROSITE profiles:**  PS50056 |  |
| Tensin-type phosphatase domain | 14-185 | dephosphorylate the D3 position of the inositol ring of PIP3 | 14 | **InterPro:** IPR029023 | ^8^ |
|  |  |  |  | **PROSITE profiles:**  PS51181 |  |
| PTEN, phosphatase domain | 24-181 | Protein phosphatase with dual specificity, acting on phosphorylated proteins | 14 | **InterPro:** IPR045101 | ^9^ |
|  |  |  |  | **CDD:**  cd14509 |  |
| Tyrosine-specific protein phosphatase, PTPase domain | 61-182 | Removal of phosphate group linked with tyrosine, using a cysteinyl-phosphate enzyme intermediate | 13 | **InterPro:** IPR000242 | ^10^ |
|  |  |  |  | **PROSITE profiles:**  PS50055 |  |
|  |  |  |  | **SMART:**  SM00194 |  |
|  |  |  |  | **PRINTS:**  PR00700 |  |
|  |  |  |  | **Pfam:**  PF00102 |  |
| Protein-tyrosine phosphatase, catalytic | 23-183 | key regulatory component in signal transduction pathways  (MAP Kinase Pathway) | 14 | **InterPro:** IPR003595 | ^7^ |
|  |  |  |  | **SMART:**  SM00404 |  |

**Table 19:** ERRAT and PROCHECK prediction of models generated by HHPred server

| Mutation | Quality Factor (ERRAT) | Residues in most favored regions | | 3D Model Assessment |
| --- | --- | --- | --- | --- |
| G129E | 54.5961 | | 88.20% | Not Appropriate |
| H123R | 49.4413 | | 89.30% | Not Appropriate |
| C124R | 61.8182 | | 86% | Not Appropriate |
| R130G | 60.0583 | | 89.80% | Not Appropriate |
| M35R | 55.6522 | | 87.60% | Not Appropriate |
| L70P | 54.4944 | | 87.80% | Not Appropriate |
| R130Q | 49.2492 | | 87.60% | Not Appropriate |
| L112P | 55.3073 | | 88.70% | Not Appropriate |
| H93R | 50 | | 89.30% | Not Appropriate |
| D252G | 56.4171 | | 88.70% | Not Appropriate |
| G132V | 59.8916 | | 89.30% | Not Appropriate |
| R173C | 52.7933 | | 88.20% | Not Appropriate |
| R173H | 56.8182 | | 87.30% | Not Appropriate |
| V119L | 60.4651 | | 88.40% | Not Appropriate |
| I135T | 54.5455 | | 88.20% | Not Appropriate |
| WILD | 53.5809 | | 89.50% | Not Appropriate |

**Table 20:** ERRAT and PROCHECK prediction of models generated by Alphafold server

| Mutation | Quality Factor (ERRAT) | Residues in most favored regions | 3D Model Assessment |
| --- | --- | --- | --- |
| G129E | 91.9355 | 84.10% | Not Appropriate |
| H123R | 89.2638 | 82.90% | Not Appropriate |
| C124R | 86.9436 | 83.70% | Not Appropriate |
| R130G | 87.346 | 82.60% | Not Appropriate |
| M35R | 89.1304 | 82.40% | Not Appropriate |
| L70P | 90.991 | 83.40% | Not Appropriate |
| R130Q | 91.7431 | 82.10% | Not Appropriate |
| L112P | 89.8773 | 82.90% | Not Appropriate |
| H93R | 87.8788 | 82.90% | Not Appropriate |
| D252G | 88.9552 | 82.90% | Not Appropriate |
| G132V | 86.7284 | 81.30% | Not Appropriate |
| R173C | 88.3436 | 82.90% | Not Appropriate |
| R173H | 89.7516 | 83.70% | Not Appropriate |
| V119L | 88.9231 | 81.50% | Not Appropriate |
| I135T | 90.7121 | 83.20% | Not Appropriate |
| WILD | 91.7178 | 81.30% | Not Appropriate |

**Table 21:** ERRAT and PROCHECK prediction of models generated by trRosetta server

| Mutation | Quality Factor (ERRAT) | Residues in most favored regions | 3D Model Assessment |
| --- | --- | --- | --- |
| G129E | 88.4718 | 85.40% | Not Appropriate |
| H123R | 91.3889 | 86.00% | Not Appropriate |
| C124R | 90.8333 | 86.80% | Not Appropriate |
| R130G | 87.2576 | 87.00% | Not Appropriate |
| M35R | 89.9729 | 87.30% | Not Appropriate |
| L70P | 88.1356 | 85.10% | Not Appropriate |
| R130Q | 90.5045 | 89.80% | Not Appropriate |
| L112P | 90.4494 | 85.40% | Not Appropriate |
| H93R | 89.7019 | 85.70% | Not Appropriate |
| D252G | 90.4192 | 89.20% | Not Appropriate |
| G132V | 86.3128 | 85.40% | Not Appropriate |
| R173C | 89.071 | 86.80% | Not Appropriate |
| R173H | 88.7052 | 86.50% | Not Appropriate |
| V119L | 88.8889 | 87.90% | Not Appropriate |
| I135T | 89.0805 | 85.40% | Not Appropriate |
| WILD | 87.7049 | 85.40% | Not Appropriate |

**Table 22:** GalaxyRefine server’s result of mutant and wild proteins of PTEN

| Mutation | GDT-HA | RMSD | MolProbity | Clash Score | Poor Rotamers | Rama favored |
| --- | --- | --- | --- | --- | --- | --- |
| G129E | 0.9628 | 0.378 | 1.644 | 9.6 | 0.5 | 97.3 |
| H123R | 0.9603 | 0.396 | 1.768 | 9.9 | 0.5 | 96.3 |
| C124R | 0.9566 | 0.403 | 1.839 | 11.8 | 1.1 | 96.5 |
| R130G | 0.9541 | 0.416 | 1.717 | 11.6 | 0 | 97.3 |
| M35R | 0.956 | 0.405 | 1.793 | 12 | 0.5 | 96.8 |
| L70P | 0.9684 | 0.374 | 1.689 | 9.9 | 0.3 | 97 |
| R130Q | 0.9584 | 0.393 | 1.764 | 11.3 | 1.1 | 97 |
| L112P | 0.9535 | 0.415 | 1.828 | 10.3 | 0.8 | 95.8 |
| H93R | 0.9615 | 0.384 | 1.555 | 10.9 | 0.5 | 98.3 |
| D252G | 0.9733 | 0.364 | 1.769 | 12.2 | 0.3 | 97 |
| G132V | 0.9498 | 0.433 | 1.797 | 13 | 0.5 | 97 |
| R173C | 0.9603 | 0.393 | 1.593 | 12 | 0.5 | 98.3 |
| R173H | 0.9566 | 0.391 | 1.748 | 10.7 | 0.3 | 96.8 |
| V119L | 0.9591 | 0.392 | 1.752 | 10.8 | 0.8 | 96.8 |
| I135T | 0.9677 | 0.371 | 1.785 | 9.8 | 0.3 | 96 |
| Wild | 0.9665 | 0.378 | 1.685 | 14.2 | 1.1 | 98 |

**Table 23:** ERRAT and PROCHECK prediction of refined by GalaxyRefine server

| Mutation | Quality Factor (ERRAT) | Residues in most favored regions | 3D Model Assessment |
| --- | --- | --- | --- |
| G129E | 91.0543 | 91.80% | Appropriate |
| H123R | 93.949 | 90.60% | Appropriate |
| C124R | 91.4013 | 90.40% | Appropriate |
| R130G | 89.3891 | 91.70% | Appropriate |
| M35R | 92.6518 | 92% | Appropriate |
| L70P | 90.8805 | 89% | Not Appropriate |
| R130Q | 89.9054 | 90.40% | Appropriate |
| L112P | 92.3077 | 89% | Not Appropriate |
| H93R | 94.5338 | 90.40% | Appropriate |
| D252G | 88.024 | 91.20% | Appropriate |
| G132V | 96.3303 | 92.60% | Appropriate |
| R173C | 93.9103 | 92.80% | Appropriate |
| R173H | 89.8734 | 89% | Not Appropriate |
| V119L | 95.5556 | 89% | Not Appropriate |
| I135T | 91.4826 | 90.60% | Appropriate |
| WILD | 90 | 93% | Appropriate |

**Table 24:** Gridbox parameter selected for docking study by AutodockVina

| SNP ID | Grid Box Parameter | | | Box size | | | Exhaustiveness |
| --- | --- | --- | --- | --- | --- | --- | --- |
|  | center_x | center_y | center_z | size_x | size_y | size_z |  |
| H123R | -26.85 | - 27.114 | -27.278 | 90 | 90 | 90 | 8 |
| C124R | -25.51 | -24.792 | -30.174 | 90 | 90 | 90 | 8 |
| R130G | -23.159 | -19.813 | -18.46 | 90 | 90 | 90 | 8 |
| M35R | -25.644 | -37.847 | -21.166 | 90 | 90 | 90 | 8 |
| L70P | -22.928 | -25.102 | -24.405 | 90 | 90 | 90 | 8 |
| D252G | -25.198 | -19.125 | -29.187 | 90 | 90 | 90 | 8 |
| R173C | -25.332 | -23.273 | -40.491 | 90 | 90 | 90 | 8 |

**Table 25:** Binding affinity and rmsd value for mutants and wildtype predicted by AutodockVina

| SNP ID | Binding Affinity (Mutant) - K/cal | Distance from rmsd (Mutant) | Binding Affinity (Wild) - K/cal | Distance from rmsd (Wild) |
| --- | --- | --- | --- | --- |
| H123R | -6.8 | 10.002 | -7.2 | 0 |
| C124R | -6.4 | 25.714 | -6.8 | 19.785 |
| R130G | -6.8 | 9.38 | -6.4 | 26.43 |
| M35R | -6.7 | 33.284 | -7.3 | 21.588 |
| L70P | -6.1 | 20.575 | -6.7 | 20.609 |
| D252G | -6.8 | 3.572 | -7.3 | 6.098 |
| R173C | -6.4 | 17.223 | -7.5 | 0 |
